# A comprehensive *Schizosaccharomyces pombe* atlas of physical transcription factor interactions with proteins and chromatin

**DOI:** 10.1101/2024.08.20.607873

**Authors:** Merle Skribbe, Charlotte Soneson, Michael B. Stadler, Michaela Schwaiger, Vishnu N. Suma Sreechakram, Vytautas Iesmantavicius, Daniel Hess, Eliza Pandini Figueiredo Moreno, Sigurd Braun, Jan Seebacher, Sebastien A. Smallwood, Marc Bühler

## Abstract

Transcription factors (TFs) are key regulators of gene expression, yet many of their targets and modes of action remain unknown. In *Schizosaccharomyces pombe*, one-third of TFs are solely homology-predicted, with few experimentally validated. We created a comprehensive library of 89 endogenously tagged *S. pombe* TFs, mapping their protein and chromatin interactions using immunoprecipitation-mass spectrometry and chromatin immunoprecipitation sequencing. Our study identified protein interactors for half the TFs, with over a quarter potentially forming stable complexes. We discovered DNA binding sites for most TFs across 2,027 unique genomic regions, revealing motifs for 38 TFs and uncovering a complex regulatory network of extensive TF cross- and autoregulation. Characterization of the largest TF family revealed conserved DNA sequence preferences but diverse binding patterns, and identified a repressive heterodimer, Ntu1/Ntu2, linked to perinuclear gene localization. Our TFexplorer webtool makes all data interactively accessible, offering new insights into TF interactions and regulatory mechanisms with broad biological relevance.

**HIGHLIGHTS:**

- Comprehensive strain library of endogenously tagged *S. pombe* TFs
- Experimentally determined atlas of TF interactions with proteins and chromatin
- TFexplorer web application for interactive exploration of TF interactomes
- Identification of repressive Nattou complex linked to perinuclear gene localization

## INTRODUCTION

Transcription factors (TFs) play a pivotal role in gene regulation, orchestrating gene expression by binding to DNA in a sequence-specific manner and recruiting effector proteins. Despite their critical role, the regulatory targets and modes of action of many TFs remain unknown. Recently, large-scale characterizations of individual TFs^1–6^ have expanded our understanding of gene regulation. Projects like ENCODE^5,6^ have substantially advanced genome-wide investigations of TF binding sites, and many physical TF interactions have been identified using proximity labeling or immunoprecipitation-mass spectrometry (IP-MS) screens^7^ and scalable binary protein-protein interaction (PPI) assays like yeast-2-hybrid (Y2H)^2–4^. However, these screens often rely on ectopic expression of TFs, do not reflect physiological conditions, or face technical challenges such as antibody comparability. The budding yeast community addressed these challenges by creating large collections of strains^8,9^ that enable researchers to study TF interactions at the systems level^10–12^. However, comparable comprehensive datasets are not available for other organisms.

The fission yeast *Schizosaccharomyces pombe* is an excellent model to study gene regulation due to its conserved regulatory processes and genetic features shared with metazoans^13^, providing insights into fundamental biological phenomena not easily studied in other model organisms. Despite its instrumental role in uncovering principles of epigenetic genome regulation and transcription, the TFs encoded by the *S. pombe* genome have received little attention. Among the 93 genes annotated with “DNA-binding transcription factor activity” on PomBase^14^ (as of March 2024), excluding general TFs like RNA polymerase II (Pol II) cofactors, approximately one-third are inferred solely from homology^14^. Moreover, only a few TFs have been extensively studied experimentally^15^.

To address this gap, we generated a strain library of 89 predicted *S. pombe* TFs endogenously tagged with a uniform epitope tag and used it to create an experimentally determined atlas of their physical interactions with proteins and chromatin in vegetatively growing cells. We found putative protein interactions for approximately half of the investigated TFs, including specific interactions with conserved co-activators and an intriguing connection to a protein with a canonical role in purine metabolism, indicating an unknown chromatin-related function. We also found abundant interactions between TFs and the regulatory phospho-binding 14-3-3 proteins, suggesting a conserved regulatory mechanism for TFs. Our genome-wide TF binding analyses revealed diverse binding patterns and identified genomic regions with a potentially unique regulatory environment characterized by a high occupancy of TFs. We uncovered a regulatory network of extensive TF cross- and autoregulation and observed potential position-dependent TF binding preferences. By providing motif information alongside TF binding and interaction data, we enable the investigation of motif selection across a comprehensive set of TFs, facilitating studies on context-dependent TF binding and cooperativity. Finally, we characterized the largest TF family, revealing conserved DNA sequence preferences and identifying a repressive heterodimer, Ntu1/Ntu2, linked to perinuclear gene localization, enabling future studies into gene repression in the context of subnuclear localization. Together, our results underscore the complex nature of TF interactions and their regulatory potential. To support further research into how TFs regulate gene expression, we provide all data, metadata, and code with detailed explanations of analysis steps and parameter choices and developed the TFexplorer web tool for interactive and user-friendly exploration of all TF interactomes.

## RESULTS

### Generation of a comprehensive *S. pombe* strain collection to establish TF interactomes

To determine the interactions of fission yeast TFs with proteins and chromatin experimentally, we generated a strain collection with each putative TF endogenously tagged with an affinity epitope (3xFLAG-tag) (Figure 1A). From the curated list of 93 putative TFs^14^, we chose to investigate 89 TFs, excluding four TFs (mat1-Mi, mat1-Mc, mat2-Pi, mat3-Mc), as they are associated with the fission yeast mating types^16^ *h^+^* and *h^-^* and our investigation is conducted in haploid, homothallic *h^-^* strains. The 89 included TFs represent diverse DNA-binding domains (DBDs) across more than 14 conserved families, as identified by the NCBI Conserved Domain Search^17^ (Figure 1B). Additionally, 18% of these TFs have predicted human orthologs^14^ (Figure 1B), and more than half possess zinc-coordinated DBDs, mirroring the proportion in the human TF catalog (807/1639 = 49%)^18^. The majority of *S. pombe* TFs are expressed in vegetatively growing cells, based on mRNA expression profiling (mRNA-seq) (Figure 1C, Figure S1A), allowing us to study them under consistent growth conditions. To determine TF-protein interactions, we performed low stringency (150 mM NaCl) IP-MS screening on all 89 TFs, followed by a higher stringency screen (500 mM NaCl) to validate putative interactions (Figure 1D, Figure S1B). As expected, TFs with very low or undetectable expression levels showed no or poor enrichment in our IPs, like Mei4, Atf31, Atf21, Cuf2, and Rsv1, which are distinctly upregulated six hours into meiosis^19^ (Figure 1C, Figure S1A). Four other TFs (Cha4, Cbf12, Sre2, and SPBC1348.12) were not identified (Figure S1A), potentially due to low expression or poor tag accessibility. To investigate TF association with chromatin, we conducted chromatin immunoprecipitation coupled to next-generation sequencing (ChIP-seq) for all 80 successfully immunopurified TFs (Figure 1E, Figure S1C). Using this strain library of TFs endogenously tagged with the same epitope tag, we established comprehensive and comparable transcription factor interactomes in vegetatively growing cells cultured under optimal conditions (30°C, rich medium).

**Figure 1.**
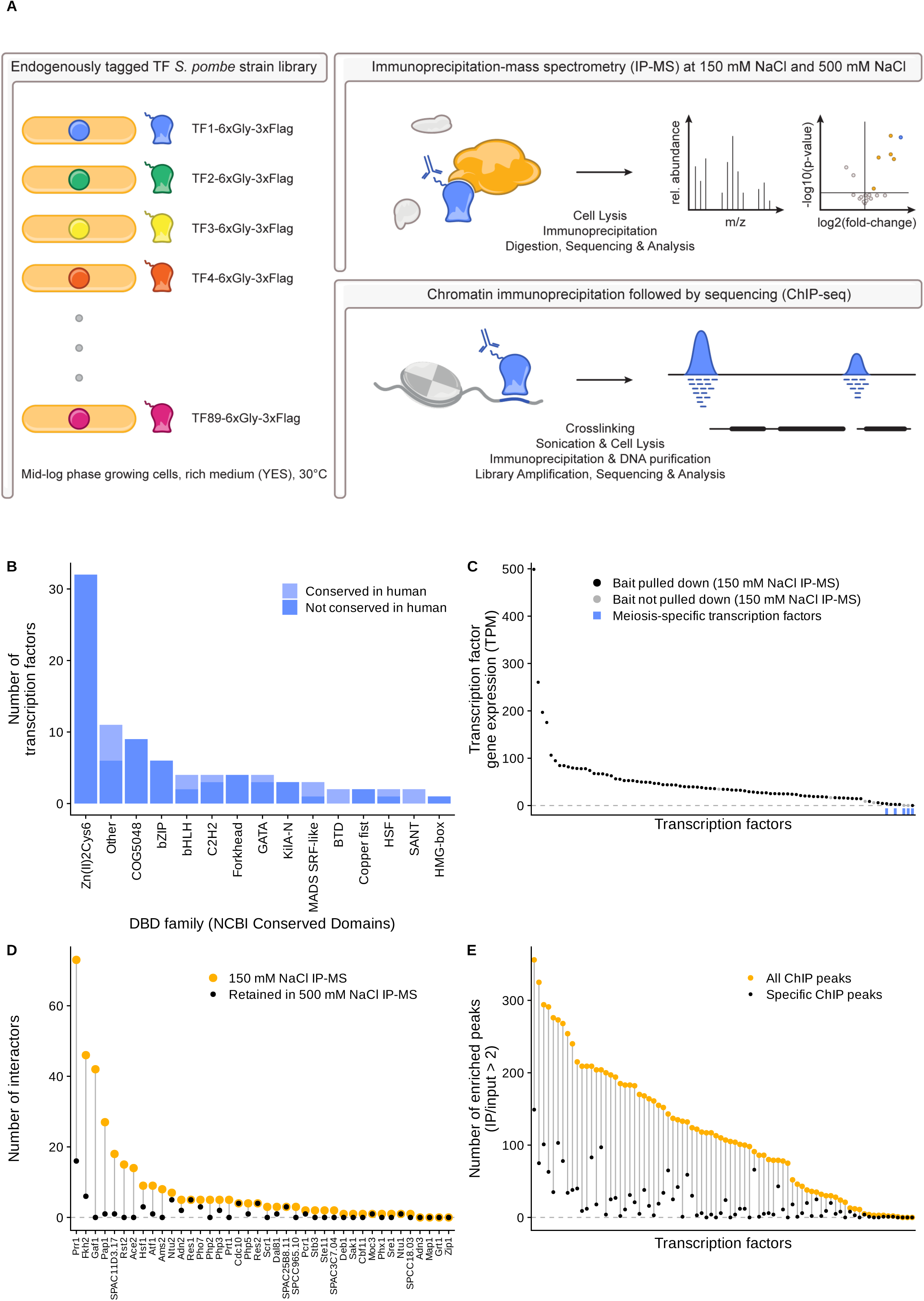
Generation of a comprehensive *S. pombe* strain collection to establish TF interactomes. (A) Schematic of the screening process to identify TF interactions with proteins and chromatin. Each strain contains one of 89 TF genes tagged at the endogenous locus with a 3xFLAG affinity epitope. Cells cultured under optimal conditions (30°C, rich medium) were subjected to IP-MS at 150 mM and 500 mM NaCl, and to ChIP-seq. (B) TF distribution across DBD families as identified by the NCBI CD-Search^17^. Lighter color indicates TFs with human orthologs^14^. (C) mRNA expression levels (wild-type) of investigated TFs, ordered by decreasing transcripts per million (TPM). Grey dots represent TFs not identified in the low salt IP-MS screen and blue bars indicate meiosis-specific TFs. (D) Number of TF interactors identified in 150 mM (orange) and retained at 500 mM NaCl (black) IP-MS screens (adj. p-value < 2e-4 & log2(fold-change) > 1). (E) Number of enriched ChIP peaks (IP/input > 2) for investigated TFs, all peaks in orange and specific peaks (bound by at most four TFs) in black. See also Figure S1.

### More than one-quarter of investigated TFs interact with other proteins under stringent conditions

We leveraged the uniform epitope tag used for all IPs to perform comparative analyses, allowing us to identify TF-specific interactions by comparing co-purifying proteins across multiple IP experiments. Each TF IP experiment, conducted in triplicates, was compared to a comprehensive control group (complement), which included all other anti-FLAG IPs and untagged controls using the same experimental protocol (tube- or plate-based 150 mM NaCl or 500 mM NaCl IP-MS protocols, see methods) (Figure S2A). Putative interactors were identified based on their significant enrichment in each TF IP-MS experiment relative to its control group by calculating a moderated t-statistic (Figure 2A, Figure 2B), which accounts for variability in protein detection^20^.

**Figure 2.**
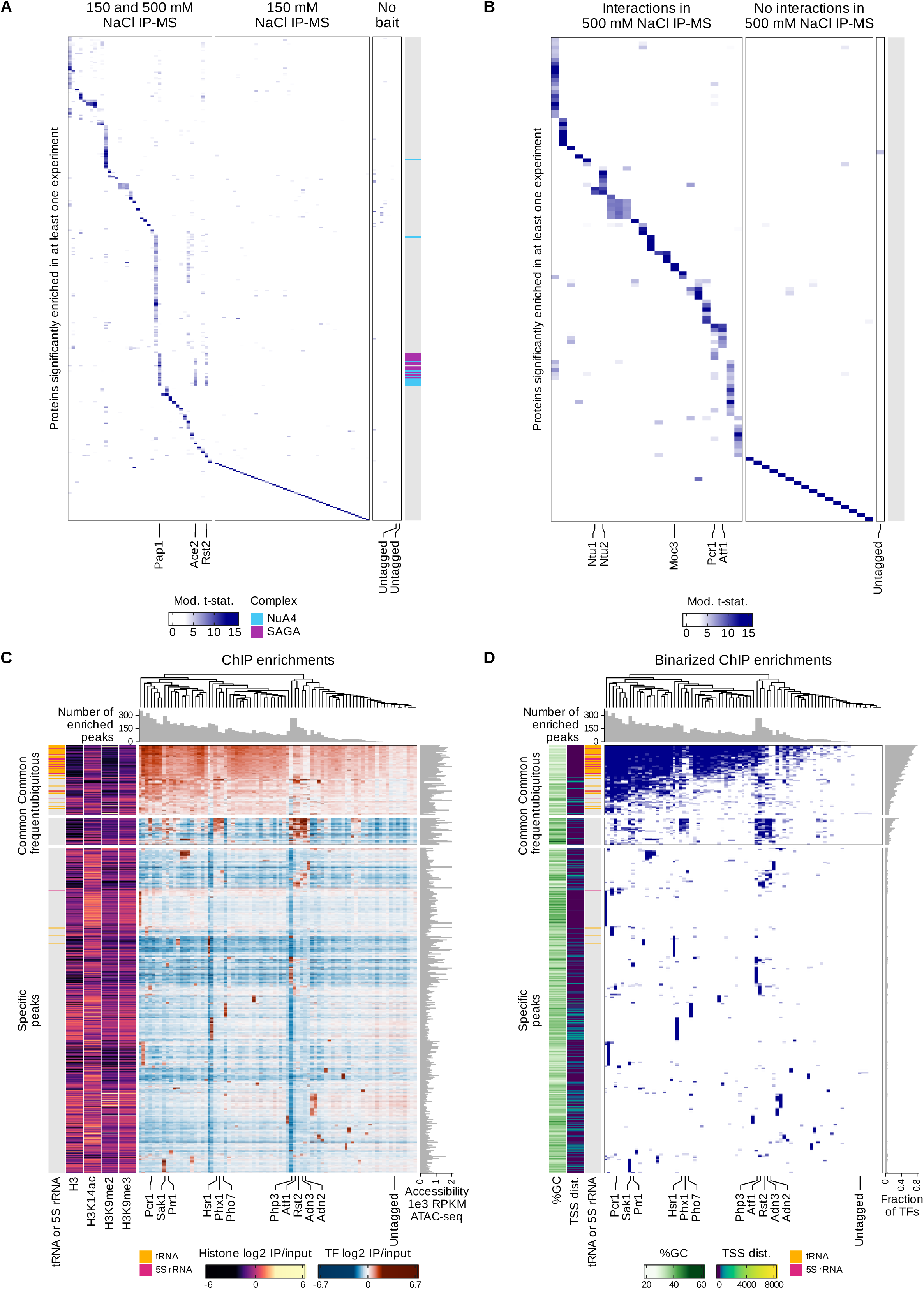
Investigated TFs exhibit abundant protein interactions and diverse genome-wide binding patterns. (A) and (B) Heatmap of moderated t-statistics for 150 mM NaCl IP-MS (A) and 500 mM NaCl IP-MS (B) screens, respectively. Proteins significantly enriched in at least one experiment (IP vs complement, adj. p-value < 2e-4 & log2(fold-change) > 1, rows) in TF and untagged control IP-MS experiments (triplicate averages, columns). Proteins (except TFs without interactions) are clustered using hierarchical clustering (complete linkage) based on the correlation between their t-statistic vectors. Columns are reordered to follow the order of TFs in rows. Complex annotations indicate NuA4 and SAGA subunits in blue and purple, respectively. Heatmap columns in (A) are split into “No bait” enrichment, “150 mM NaCl IP-MS” (TFs only investigated in the low salt screen), and “150 and 500 mM NaCl IP-MS” (TFs additionally investigated in the high salt screen). Heatmap columns in (B) are split into untagged control, “No interactions in 500 mM NaCl IP-MS” (TFs with no detected interactors at high salt), and “Interactions in 500 mM NaCl IP-MS” (TFs with interactions at high salt). (C) Heatmap colored by ChIP enrichment values (log2(IP/input)). Heatmap peaks (rows) are split into specific peaks (bound by at most four TFs) and common peaks, which are further subdivided into frequent and ubiquitous peaks by k-means clustering of their enrichment values across TF ChIPs. Annotations include enriched peaks per TF (top), accessibility data^104^ (ATAC-seq, right), presence of tRNA (orange) or 5S rRNA (pink) genes (left), and ChIP-seq data (left) for H3/H3K14ac^105^ and H3K9me2/H3K9me3^106^. (D) Heatmap as in (C) showing binarized ChIP enrichments. Binary values (blue: TRUE, white: FALSE) indicate average ChIP enrichment values of log2(IP/input) > 1. Additional annotations include fraction of TFs enriched per region (right), and GC-content (%) and distance to TSS (left). See also Figure S2.

With our primary goal of highlighting the most important findings, we report interactions using a stricter significance cutoff than is typically employed (adjusted p-value < 2e-4 & log2(fold-change) > 1), unless specified otherwise. However, all data are available in the TFexplorer webtool with user-definable p-value cutoffs, enabling further exploration with adjustable stringency (show in [TFexplorer](https://data.fmi.ch/TFexplorer/pombe.html#IPMS)). Based on the low salt IP-MS data, we classified each TF into one of three categories: “No bait” enrichment, “150 mM NaCl IP-MS” for TFs subjected only to the low salt IP-MS screen, and “150 and 500 mM NaCl IP-MS” for TFs also subjected to the high salt IP-MS screen to assess interaction stability (Figure 2A). We identified interactions for 43 TFs, yielding a total of 352 interactions in the low salt screen. In the high salt screen, we observed 110 interactions for 24 TFs. For comparison, using a lenient cutoff (adjusted p-value < 0.01 & log2(fold-change) > 1), we found 881 interactions for 60 TFs in the low salt screen and 202 interactions for 29 TFs in the high salt screen (show in [Report](https://fmicompbio.github.io/Spombe_TFome/ms_figure_5_ipms.html#Facts_and_figures)).

These results indicate that under optimal growth conditions, approximately half of the investigated TFs potentially interact with other proteins, with more than one-quarter possibly forming stable interactions.

### Genome-wide analysis of TF binding reveals diverse patterns and high occupancy target regions

To determine genome-wide TF binding sites, we conducted ChIP-seq on the 80 successfully immunopurified TFs, performed in duplicates, and called peaks individually on pooled replicates. Retaining peaks with at least 1.75-fold enrichment in at least two samples and excluding blacklisted regions (see methods), we identified 2,027 unique peak regions across all experiments. We identified ChIP peaks with at least 2-fold enrichment (IP over input) for 77 TFs (Figure 1E, Figure S1C), resulting in a total of 9,365 peaks. Peak enrichments were strongly correlated between replicates, demonstrating the quality and reproducibility of our ChIP-seq data (Figure S2B). The number of enriched peaks per TF ranged from 1 to 356 (Figure 1E, Figure S1C), with 50% (interquartile range) of TFs identifying between 38 and 183 peaks. TFs exhibited diverse binding patterns (Figure 2C, Figure 2D), ranging from singular to multiple hundred binding sites. Most TF binding sites reside in more accessible regions indicated by low levels of H3, and many peaks are characterized by elevated H3K14ac levels (Figure 2C).

We estimated the number of TF-bound gene promoters, defined as a 1 kb bin centered on the transcription start site (TSS), and found that 32% (2,347) of all gene promoters and 27% (1,400) of protein-coding gene promoters are associated with TFs (show in [Report](https://fmicompbio.github.io/Spombe_TFome/ms_figure_3_chip.html#Facts_and_numbers_-_ChIP-seq)). Highly expressed protein-coding genes, as measured by mRNA-seq, tend to be bound by at least one TF (Figure S2C). In contrast, 68% (4,922) of all gene promoters show no TF enrichment. Comparing bound regions (2,027) across all TF ChIP experiments, we observed striking disparities in the number of TFs binding to a given site, ranging from a single TF to 65 (>80%) (Figure 2D). We leveraged the comparability of all TF ChIPs to classify all identified peak regions based on the number of TFs that bind to them. We distinguished “specific” peaks, detected in less than 5% of TF ChIP experiments (max. overlap of four TFs), from “common” peaks, detected in more than 5% of experiments (Figure 2C, Figure 2D). This cutoff accounts for specific binding sites of known trimeric TF complexes, such as the CCAAT-binding factor^21^ and MBF complex^22^.

The common peak regions in our ChIP-seq dataset resemble high occupancy target (HOT) regions observed in large-scale TF ChIP-seq datasets across multiple organisms^5,6,23,24^. Initially dismissed as artifacts, HOT regions are increasingly recognized as genuine biological phenomena, yet remain poorly understood^25–27^. Further examining these regions, we found a strong enrichment for tRNA and 5S rRNA genes specific to common peak regions with TF ChIP peaks overlapping these genes showing moderate, uniform enrichments (Figure 2C). An unsupervised k-means clustering approach based on peak enrichments (IP/input of replicate averages) supported this observation, clustering them into a group that we named “common ubiquitous” peaks. Notably, ChIP samples exhibited a GC-bias with more reads in GC-rich regions compared to AT-rich ones. Because this trend was consistent, we could correct for this bias (see methods), but it did not affect these weak enrichments observed in common ubiquitous peaks. Given the unlikely specific binding of so many TFs to these Pol III-transcribed genes and that peak enrichments are uniform across most ChIPs only at these loci (Figure 2C), we consider “common ubiquitous” peaks technical artifacts. Their absence in the untagged control suggests that this bias originates during bait purification, regardless of whether the tagged TF binds to DNA. Unlike “common ubiquitous” peaks, the second cluster, termed “common frequent” peaks, lacks a singular defining feature. Peaks in this cluster are detected for 5 to 26 TFs (Figure 2D), with an average of 9.2 TFs per region. They are predominantly nucleosome-free and accessible (Figure 2C, Figure 2D), and, like specific peak regions, they have distinct peak enrichments and average GC-content. We consider these binding events true and conclude that these “common frequent” regions are *S. pombe* HOT regions, similar to previously described HOT regions in other organisms^23,25,27^. The number of TFs detected in HOT regions varies, but most of these regions contain a core set of bound TFs (Php3, Sak1, Pcr1, Prr1, Atf1, Rst2, Adn2, Adn3, Hsr1, Phx1, and Pho7) (Figure 2C). Some studies suggest that HOT regions arise from indirect binding and extensive multivalent and weak TF-TF interactions^27–29^, potentially enabled by their intrinsically disordered regions (IDRs). Consequently, the observed co-occupancy could be attributed to indirect binding events rather than individual, direct TF-DNA contacts. However, our IP-MS dataset did not reveal a clear difference in interaction patterns between HOT TFs and all other TFs (Figure S2D), though given that IDR-driven interactions are likely weak, the absence of increased interactions measured by IP-MS might be expected.

In summary, our comprehensive ChIP-seq analysis of 80 TFs revealed diverse TF binding patterns, with approximately one-third of gene promoters bound by at least one TF. We identified 247 common ubiquitous regions likely attributable to technical artifacts, 94 genuine HOT regions, and 1,686 specific peak regions, thereby advancing our understanding of TF interactions with the *S. pombe* genome.

### Extensive cross- and autoregulation among *S. pombe* TFs

In addition to investigating genome-wide TF binding events, we analyzed the regulatory network among TFs. Examining the promoters of all 89 TFs, we discovered that 43 TF promoters are bound by at least one other TF, with 26 promoters bound by multiple TFs (Figure 3A). Particularly, Atf1, a key regulator of the *S. pombe* stress response^30–33^, is highly connected (Figure 3B). Atf1 binds its own promoter and the promoters of ten other TFs. In turn, the *atf1^+^*promoter is bound by eleven TFs, including Atf1 itself and its heterodimer partner Pcr1. Both Atf1 and Pcr1 also bind to the *pcr1^+^* promoter, indicating that they co-regulate their own genes (this study and Eshaghi *et al.*^34^).

**Figure 3.**
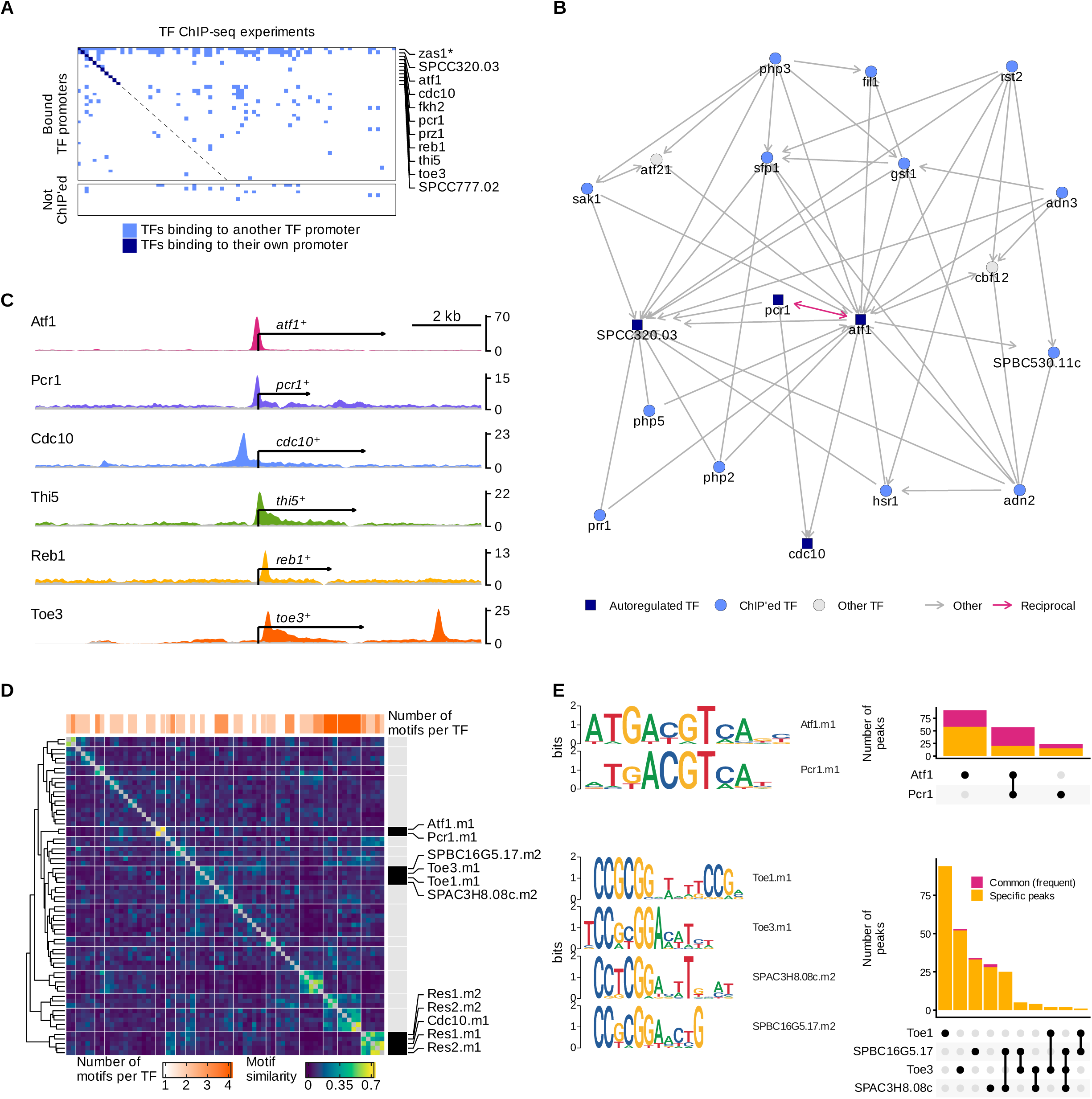
Analysis of TF regulatory networks and *de novo* DNA-binding motif identification. (A) Heatmap showing binarized ChIP enrichments at TF gene promoters (1 kb bin centered on TSS) (rows). Binary values (blue: TRUE, white: FALSE) indicate average ChIP enrichment values of log2(IP/input) > 1 with dark blue further indicating TF ChIP enrichments over their own promoters. Heatmap rows are split into bound TF gene promoters of investigated TFs, and all promoters of non-investigated TFs. Diagonal (dashed line) matches each TF ChIP with its promoter. Asterisk (*) indicates tRNA gene within *zas1^+^* promoter (considered artifact detected in 48 TF ChIPs). (B) Atf1-centric network representation of (A) with nodes representing TFs: dark blue for autoregulated, light blue for ChIP’ed, and grey for not investigated (other). Edges indicate detected promoter binding events, starting at the binding TF and ending at the promoter of the bound TF. Only edges connecting to Atf1 or other TFs connecting to Atf1 are shown. The Zas1 node was excluded based on (A). The full network is available in the [Report](https://data.fmi.ch/TFexplorer/pombe.html?tf=Esc1,SPAC3F10.12c#IPMS). (C) Genome browser views of ChIP (colored) and input (grey) fragment densities for six TFs around their gene loci. Scale: Counts per million (CPM) per base-pair (bp), smoothed over 101 bp windows using a running mean. All genes are oriented to align at their TSS. Coverage dips at 3’ ends correspond to affinity tag insertion sites. (D) Heatmap of pairwise motif similarities (PCC), clustered into 17 groups. Three highlighted clusters include motif labels. Top annotation shows the number of predicted motifs per TF. Fully annotated heatmap in Figure S3B. (E) Aligned sequence logos for motifs from two highlighted clusters in (E). UpSet plots show the number of specific (orange) and common frequent (pink) ChIP peaks shared among TFs with similar motifs. See also Figure S3.

Beyond Atf1 and Pcr1, nine additional TFs bind their own promoters, suggesting autoregulation (Figure 3C, Figure S3A). Of all eleven TFs found binding to their own promoters, six have been previously shown or suggested to do so (Atf1, Pcr1, Toe3, Cdc10, Prz1, and Fkh2)^34–39^. Interestingly, certain TFs, such as Thi5, Reb1, and Toe3, bind their promoters at or downstream of their annotated TSS (Figure 3C). Though these TFs may utilize alternative TSSs, they may also regulate their genes in a position-dependent manner. A recent preprint reported regulatory regions downstream of the TSS with TF family-specific binding preferences in plants^40^ and a study of human TF binding motifs found highly preferential positioning relative to the TSS, determining TF function^41^.

Our analysis of the fission yeast TF regulatory network revealed extensive cross-regulation and suggests autoregulation for 14% of investigated TFs under optimal conditions. Additionally, our data indicate potential position-dependent TF binding preferences. Systematically evaluating if this is specific to the TF, the TF family, or the regulated genes will provide further insights into the fission yeast regulatory landscape.

### *De novo* DNA-binding motif identification indicates context-dependent TF binding

For a deeper understanding of TF binding sites, we aimed to identify DNA motifs from bound genomic regions. Initially including sequences from all peak categories, predictions were skewed towards consensus sequences in tRNA and 5S rRNA gene promoters. Only after excluding common peaks we could reliably predict individual DNA-binding motifs, further suggesting that these observed binding events might be artifactual. We then curated a list of the most confident DNA-binding motifs (see methods), yielding 67 predicted motifs for 38 TFs. Using the Pearson correlation coefficient (PCC) as a similarity metric, we identified both known and novel motif similarities and detected any duplicate motifs for the same TF, exemplified by Res1 and Res2 (Figure 3D, Figure S3B). After removing redundancies, we identified 45 unique motifs for 38 TFs. As expected, highly similar motifs were observed among members of known TF complexes, such as the MBF complex^36,42^ and the Atf1/Pcr1 heterodimer^30,43,44^ (Figure 3D). Consistent with prior work, Atf1 and Pcr1 recognize the same motif but can bind DNA independently^45^. We found them overlapping at approximately one-third of all their identified binding sites, excluding common ubiquitous peaks (Figure 3E). In contrast, we identified a group of TFs of the same DBD family, binuclear zinc cluster TFs Toe1, Toe3, SPAC3H8.08c, and SPBC16G5.17, with similar motifs but limited overlap in DNA binding sites (Figure 3D, Figure 3E). This observation shows that features beyond DNA sequence specificity determine genome binding, which became even more evident when we evaluated the proportion of available predicted motifs in the genome that TFs would bind. The percentage varied from 0.3% to 17.5%, with 50% (interquartile range) of TFs binding 1.2% to 4.5% of their available motifs (Figure S3C). For instance, Atf1 bound 7.7% of its predicted sites, consistent with previous studies^34^.

These findings underscore the importance of factors beyond sequence motifs in TF binding, such as the protein sequence context outside the DBD^46,47^ and the local chromatin environment^48^. By providing motif information in combination with TF binding and interaction data, we enable the investigation of motif selection across a comprehensive set of TFs to study context-dependent TF binding and cooperativity on a global scale or at individual loci.

### Systematic evaluation of binuclear zinc cluster TFs reveals conserved DNA sequence preferences and the Nattou heterodimer

Binuclear zinc clusters (Zn(II)_2_Cys_6_) are fungal-specific DBDs, and TFs of this family can bind to DNA as monomers, homodimers, or heterodimers, with each monomer interacting with a CGG trinucleotide or variations of it^49^. A prominent member of this family is the *Saccharomyces cerevisiae* TF GAL4, which binds to the DNA motif CGG- N_11_-CCG as a homodimer^50,51^. Our findings of highly similar, CGG-rich motifs for the binuclear zinc cluster TFs Toe1, Toe3, SPAC3H8.08c, and SPBC16G5.17, indicate a similar binding preference for *S. pombe* TFs of this family (Figure 3E). Notably, Toe3, SPAC3H8.08c, and SPBC16G5.17 had not previously been associated with specific motifs, whereas the DNA-binding motif for Toe1 is known^35^.

For a systematic evaluation of nucleotide binding preferences among binuclear zinc cluster TFs, we conducted an unbiased k-mer enrichment analysis. This entailed comparing the frequency of any k-mer (DNA sequence of length k) in a TF’s specific peaks against their frequency in the control sequences (specific peak sequences of all other TFs). For instance, a 6-mer analysis of Toe3 revealed high enrichments for CCRYGG sequences (Figure 4A), matching the motif Toe1.m1 found in our previous motif identification (Figure 3E). Given that monomers interact with single trinucleotides, we performed a 3-mer enrichment analysis for any family member with at least three specific peaks (20 TFs). As a result, we found enrichments for GC-rich trimers for most TFs, with a notable preference for CGGs in various orientations (Figure 4B). Toe2, in particular, exhibited a strong preference for CGG, which was confirmed as a dual repeat motif in the *de novo* motif identification (Figure 4C), akin to the motifs discussed before. However, while motifs for Toe1, Toe3, and SPBC16G5.17 comprise everted repeats (CCGCGG), the Toe2 motif consists of inverted repeats (CGGCCG), explaining their lack of similarity. The presence of repeat-containing motifs suggests that these TFs may bind as dimers, as proposed for TFs of this family in other fungi^49^. Therefore, we searched our interactome data for family-wide TF-TF interactions, which revealed a single reciprocal interaction between two previously uncharacterized binuclear zinc cluster TFs, SPBC16G5.16 (Ntu1) and SPBC530.08 (Ntu2) (Figure 5A, Figure 5B). We observed this interaction, previously detected in a proteome-wide Y2H screen when SPBC16G5.16 served as bait^52^, reciprocally and under high stringency conditions (Figure 5C). Therefore, we propose that these proteins form a TF heterodimer complex, which we are naming the Nattou complex, with the subunits Ntu1 (SPBC16G5.16) and Ntu2 (SPBC530.08). Conversely, the observation of limited interactions amongst binuclear zinc cluster TFs implies that Toe1, Toe2, Toe3, and SPBC16G5.17 may bind to CGG repeat-containing sequences as homodimers.

**Figure 4.**
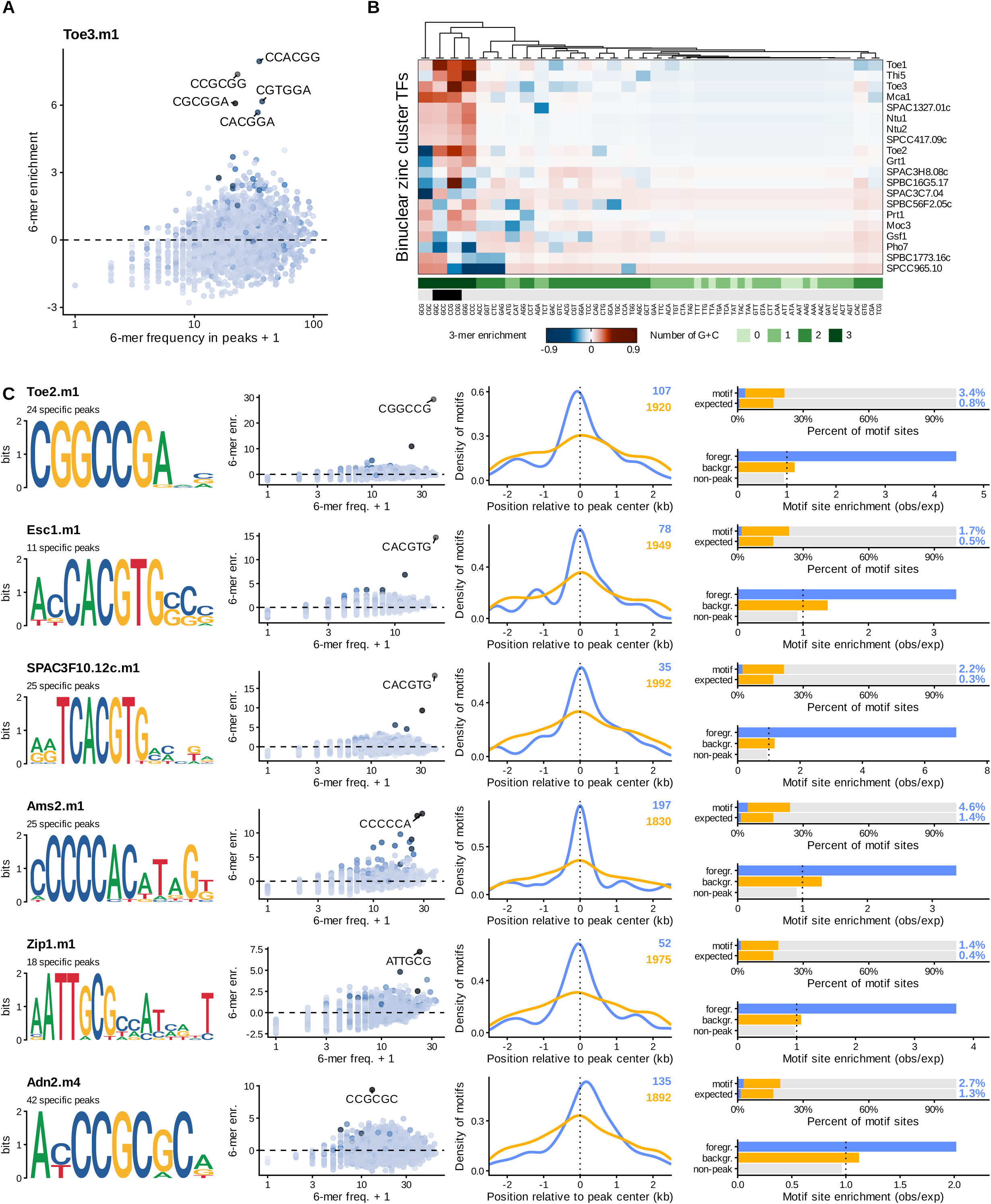
Systematic evaluation reveals conserved DNA sequence preferences for binuclear zinc cluster TFs and E-box motifs for bHLH TFs. (A) Frequency and enrichment of 6-mer DNA sequences in specific Toe3 ChIP peaks, colored by similarity to the predicted motif Toe3.m1. Darker colors indicate higher similarity. (B) Clustered heatmap of 3-mer DNA sequence enrichments in specific ChIP peaks of 20 binuclear zinc cluster TFs. GC-content in green, CGG 3-mers in any orientation in black. (C) Characterization of six predicted DNA-binding motifs. Plots (left to right): Sequence logo of the predicted motif, frequency and enrichment of 6-mers as in (A), motif location relative to peak center in foreground (all enriched TF ChIP peaks, blue) and background peaks (peaks from all other TF ChIPs, yellow), percent of motif sites observed or expected in foreground, background, and non-peak regions (grey) (top), and enrichment of observed over expected (bottom). See also Figure S4.

**Figure 5.**
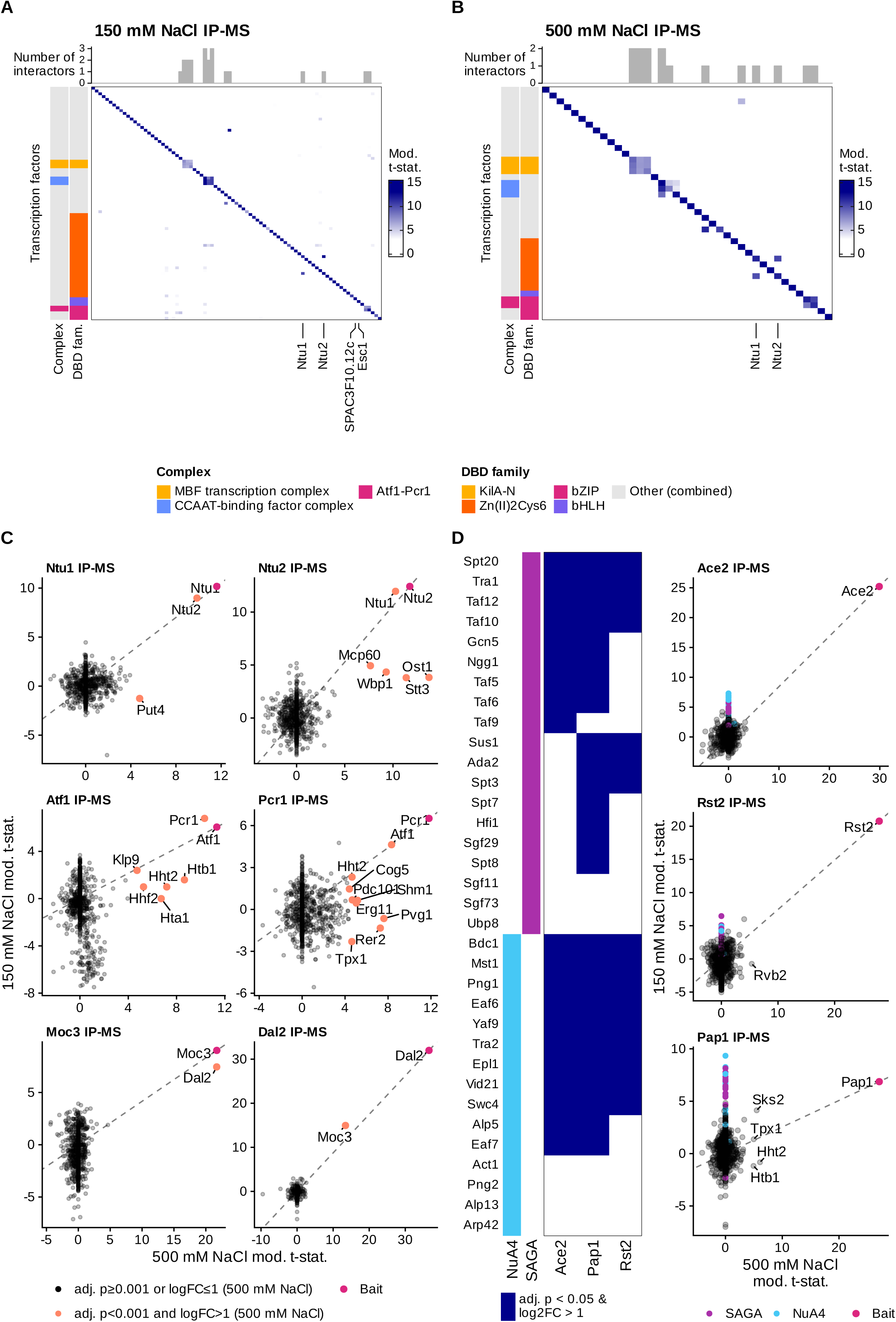
Diverse TF interactions with other TFs, coactivator complexes, histones, and a putative allantoicase. (A) and (B) TF-specific subset of moderated t-statistic result heatmaps for 150 and 500 mM NaCl IP-MS screens, respectively (IP vs complement). Negative values set to 0. TFs identified in each screen (rows) plotted for TF IP-MS experiments (triplicate averages, columns). Top annotations indicate the number of TF interactors identified for each TF (adj. p-value < 2e-4 & log2(fold-change) > 1). Row annotations indicate known TF complexes and their DBD family. (B) Scatter plots correlating moderated t-statistics from 500 and 150 mM NaCl IP-MS screens for indicated TFs (IP vs ‘high salt complement’, see methods, for all TFs except Dal2: IP vs untagged control). Proteins significantly enriched in the high salt screen (adj. p-value < 0.001 & log2(fold-change) > 1) in orange and bait in pink. Dashed line intersects bait and origin. (C) Scatter plots correlating moderated t-statistics from 500 and 150 mM NaCl IP-MS screens for Ace2, Rst2, and Pap1. SAGA and NuA4 subunits in purple and blue, respectively, and bait in pink. Dashed line intersects bait and origin. Binary heatmap (left) shows significantly enriched SAGA and NuA4 subunits (adj. p-value < 0.05 & log2(fold-change) > 1) with TRUE values in blue. See also Figure S5.

In summary, we identified novel DNA-binding motifs for Toe2, Toe3, SPAC3H8.08c, and SPBC16G5.17, and demonstrate a strong CGG trinucleotide binding preference for most *S. pombe* binuclear zinc cluster TFs. Our findings also uncovered a potential heterodimer, Ntu1/Ntu2, and provide the foundation for further investigation into the homodimerization potential of these TFs, supported by CGG repeat-containing DNA-binding motifs predicted from our ChIP-seq data.

### Basic helix-loop-helix TFs Esc1 and SPAC3F10.12c recognize near-identical E-box motifs

Besides the novel motifs we discovered for the binuclear zinc cluster TFs, we find additional DNA motifs that are particularly intriguing, prompting us to characterize them further (Figure 4C). Like the previous analysis (Figure 4A), we complemented the *de novo* motif identification, displayed as a sequence logo, with a 6-mer enrichment analysis. Then, we assessed the motif distribution, expecting the highest enrichment around the peak center. Finally, to evaluate if the motif is specific to the TF-bound regions in the genome, we counted its occurrence in enriched peaks (foreground), non-enriched peaks from all other TFs (background), and the rest of the genome (non-peak) and calculated the enrichment of observed motifs over what would be expected from a random distribution. Among the newly identified motifs, we found identical E-box motifs (CACGTG) for two *S. pombe* TFs, Esc1 and SPAC3F10.12c (Figure 4C), representing the consensus sequence for basic helix-loop-helix (bHLH) TFs^53^. Akin to the human TFs Myc, Max, and Mad, these two belong to the phylogenetic class B of bHLH TFs^54,55^ and exhibit class-typical characteristics, including CAC half-site preferences. bHLH TFs typically form dimers, with each monomer binding a CAN half-site, and the specific combination of dimerization partners influencing TF sequence specificity and function^53^. The perfectly palindromic E-box sequences (Figure 4C) suggest the possibility of homodimerization or heterodimerization within the same class. The absence of overlapping binding sites between Esc1 and SPAC3F10.12c (Figure S4A), despite their near-identical DNA-binding motifs, indicates that these TFs likely form homodimers. This hypothesis is further supported by our IP-MS data (Figure 5A) (show in [TFexplorer](https://data.fmi.ch/TFexplorer/pombe.html?tf=Esc1,SPAC3F10.12c#IPMS)) that did not indicate heterodimerization. The strong preference for a 5’ thymine in the E-box flanking sequence of the SPAC3F10.12c motif is a possible explanation for their unique binding sites.

In summary, our integrated ChIP-seq and IP-MS analyses identified E-box DNA-binding motifs for the bHLH TFs Esc1 and SPAC3F10.12c and predict their homodimerization, providing key insights to further dissect their mechanisms.

### Approximately one in three TFs engage in TF-TF interactions, with stable interactions typically formed within the same DBD family

Large-scale PPI screens in various organisms have shown a wide range of predictions for TF-TF interactions. Studies in *Drosophila melanogaster*, human, and mouse suggest that over two-thirds of investigated TFs interact with at least one other TF or themselves^1,3,7^. In contrast, a Y2H study with *Caenorhabditis elegans* proteins, which investigated nearly 90% of predicted TFs, found that only about 19% interact with themselves or other TFs^2^. To estimate the number of TF-TF interactions in *S. pombe* during optimal growth, we expanded our search beyond the binuclear zinc cluster and bHLH families, using the same strict cutoff as previously described (Figure 5A, Figure 5B). We identified 21 interactions involving 17 TFs in the low salt and 15 interactions involving 14 TFs in the high salt IP-MS screen (Figure S5A). Among these was an intriguing interaction between two LIS1 homology (LisH) domain TFs, Adn2 and Adn3, and the transcriptional coregulator Ldb1/Adn1 (Figure S5B). Ldb1/Adn1, Adn2, and Adn3 are known activators of flocculation, a process where cells rapidly aggregate in response to environmental stress. Deletion of these genes individually leads to a cell adhesion-defective phenotype^56,57^. Thus, our data suggests that their previously established functional relationship and phenotypic similarity could be due to these TFs forming a stable complex, which becomes dysfunctional upon loss of any component. Overall, we observed TF-TF interactions both within the same DBD family and across different DBD families to an equal extent, with stable complexes mostly formed between TFs of the same family (Figure 5A, Figure 5B, Figure S5A). This contrasts with findings in *D. melanogaster* and *Arabidopsis thaliana*, where extensive interactions were observed between different TF families, although not under physiological conditions^3,4^. Lowering our cutoff for qualifying interactions reveals additional (potentially transient) TF-TF interactions also in *S. pombe*, mostly between different DBD families. Using a lenient cutoff (adjusted p-value < 0.01 & log2(fold-change) > 1), we identified 39 interactions (18 bilateral) involving 30 TFs under low salt conditions, and 17 interactions (14 bilateral) involving 14 TFs under high salt conditions (Figure S5A).

Based on these findings, we estimate that approximately one-third (27-36%) of all examined *S. pombe* TFs interact with at least one other TF under optimal growth conditions. We observed that most interactions occurring within TF families lead to stable complexes, including interactions between Adn2, Adn3, and Ldb1/Adn1, which provides an explanation for their known functional relationship.

### *S. pombe* basic leucine zipper TFs might have pioneering activity

In our study, we identified TFs that specifically interact with core histone subunits of the canonical nucleosome. Although histones were abundantly pulled down in all low stringency IPs, including the untagged control, this interaction was uniquely preserved for Pap1 and the heterodimeric TFs Atf1 and Pcr1 amongst TFs investigated under high stringency conditions (Figure 5C, Figure S5C). This finding is particularly intriguing given recent structural studies that have highlighted direct interactions between certain pioneer TFs and the nucleosome core^58–60^. Pioneer TFs can bind to DNA and access their binding sites despite the presence of a nucleosome. Whereas these TFs can induce changes in the local chromatin landscape, the exact definition of pioneering TFs remains a topic of debate^60,61^.

Atf1 and Pcr1 have been implicated in various nucleosome-related processes, including the creation and maintenance of nucleosome-depleted regions^62–65^, meiosis-specific chromatin remodeling at the M26 hotspot^66,67^, and heterochromatin initiation and maintenance at the mating type locus^68–71^. Whereas Pap1 has not been linked to nucleosome remodeling, its human orthologs, Jun/Fos, have been shown to bind nucleosomes *in vitro*^72^ and associate with nucleosome-rich regions *in vivo*, potentially enhancing accessibility by recruiting chromatin remodelers^73^. Interestingly, Atf1, Pcr1, and Pap1 are among the six basic leucine zipper (bZIP) TFs in *S. pombe*^14^. The fourth bZIP TF expressed in mitotically growing cells, Zip1, did not co-purify histones under high stringency conditions (Figure S5C). The remaining two bZIP TFs, Atf21 and Atf31, are exclusively expressed during meiosis^19^ and investigating them in meiotic cells may reveal whether nucleosome binding is a characteristic shared by *S. pombe* bZIP TFs. Our finding that Pap1, Atf1, and Pcr1 co-purify with histones under stringent conditions indicates interactions with the nucleosome and suggests pioneering activity for these TFs that may extend to an entire family of *S. pombe* TFs.

### TF interactions with the conserved acetyltransferase complexes SAGA and NuA4, and a putative allantoicase

Investigating our interactome data to identify TF interactions with chromatin modifiers, we confirmed the recently characterized stable interaction between the forkhead TF Fkh2 and the Clr6 histone deacetylase complex^74,75^ (show in [TFexplorer](https://data.fmi.ch/TFexplorer/pombe.html?tf=Fkh2#IPMS)). Additionally, we discovered previously unknown interactions with co-activator complexes: three TFs (Rst2, Pap1, and Ace2) co-purified with multiple subunits of the conserved acetyltransferase complexes SAGA and NuA4 in the low salt screen (Figure 5D). Supporting the validity of these interactions, Rst2 has previously been implicated in recruiting SAGA based on genetic evidence^76^. Furthermore, we observed an interaction between the binuclear zinc cluster TF Moc3 and the putative allantoicase Dal2, previously suggested by a Y2H assay^52^ (Figure 5C). By immunopurifying Dal2, we confirmed that this interaction is detected reciprocally under high stringency conditions (Figure 5C). This finding is intriguing given the canonical role of allantoicase enzymes (Enzyme Commission number 3.5.3.4) in purine metabolism, primarily in utilizing purines as a secondary nitrogen source. The interaction between a TF and a predicted allantoicase hints at a non-canonical function for one of these proteins. Given that Moc3 is constitutively nuclear and functionally dependent on its DBD^77^, which is supported by our ChIP data showing its association with chromatin (show in [TFexplorer](https://data.fmi.ch/TFexplorer/pombe.html?tf=Moc3#ChIP-Seq)), we propose that Dal2, rather than Moc3, might have a moonlighting function.

Together, our interactome data revealed specific interactions of TFs with the conserved co-activators SAGA and NuA4, as well as with a putative allantoicase, suggesting potential roles in chromatin modification and uncovering a possible novel chromatin-related activity.

### Extensive interactions between TFs and Rad24/Rad25 suggest widespread TF regulation by 14-3-3 proteins

14-3-3 proteins are phosphoprotein-binding factors that are well-conserved among eukaryotes, forming homo-and heterodimers to modulate various cellular processes by interacting with diverse proteins, including TFs^78^. *S. pombe* has two 14-3-3 protein paralogs, Rad24 and Rad25^79^. Our study revealed that Pho7, a TF activated by phosphate starvation^80,81^, specifically interacts with Rad24 and Rad25 also under stringent conditions (Figure 6A). Notably, Rad24 is known to negatively regulate Pho7-dependent *pho1^+^* expression, with its deletion resulting in increased *pho1^+^* levels even under phosphate-replete conditions^82–84^. Mechanistically, Rad24 has been linked to the regulation of an upstream long non-coding RNA (lncRNA), which is known to interfere with *pho1^+^* expression in the absence of stress^84,85^. Our data suggests an alternative mechanism where Rad24, and potentially Rad25, directly interact with and negatively regulate Pho7, which would explain the de-repression of *pho1^+^* in *rad24^Δ^*mutants. We identified two optimal 14-3-3 binding motifs^78^ in Pho7, corresponding to phosphorylated serine and threonine residues (RVCSAP (pS230) and RSFTNP (pT463))^86–89^ (Figure 6B). These motifs flank the Pho7 DBD, indicating that Rad24/Rad25 interactions could interfere with Pho7 DNA binding, similar to the mechanism proposed for the mammalian TF FOXO4^90^.

**Figure 6.**
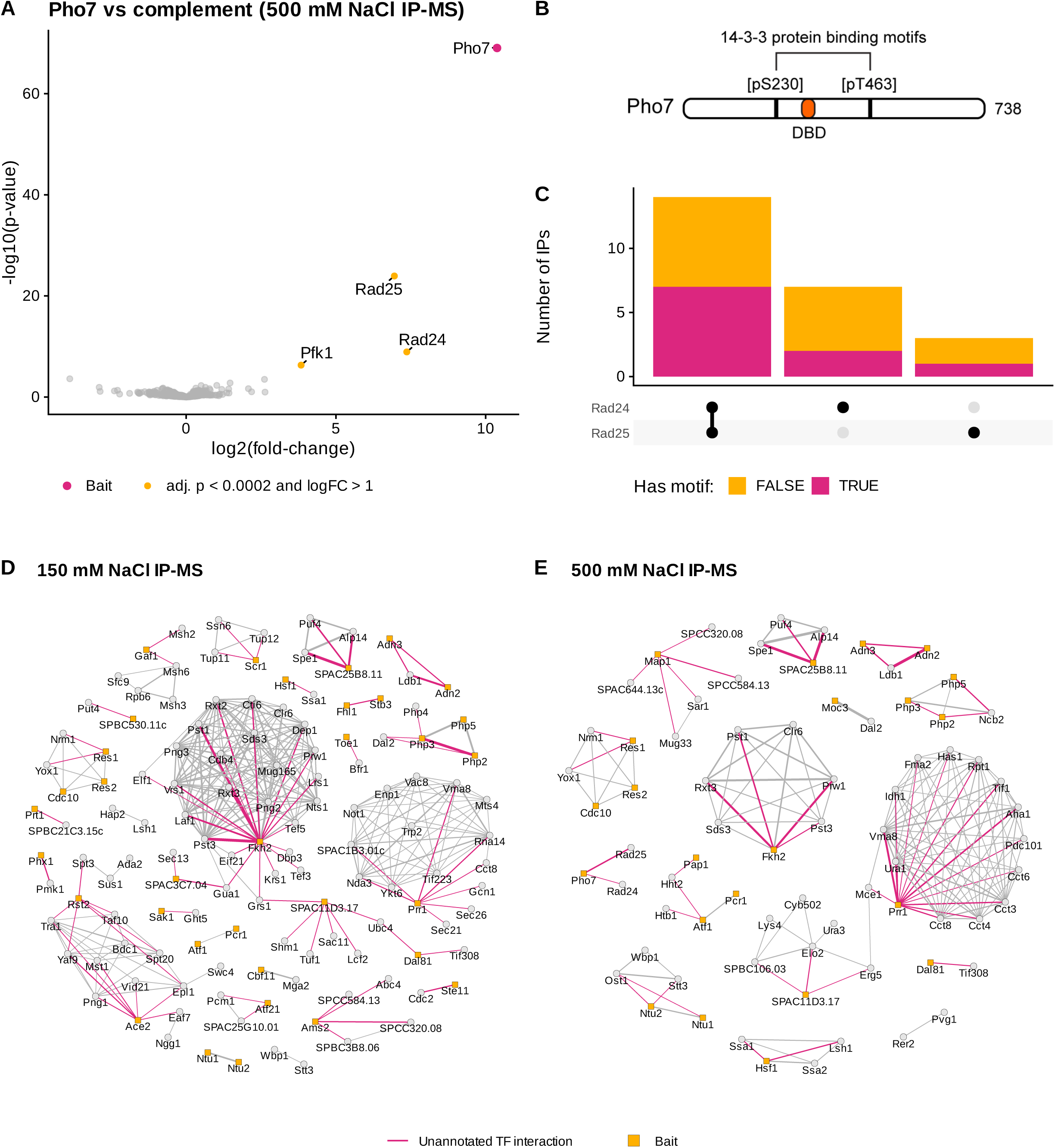
TF interaction with 14-3-3 proteins and interactively explorable networks to display putative prey-prey interactions. (A) Volcano plot for 500 mM NaCl IP-MS of Pho7 vs complement with significantly enriched proteins (adj. p-value < 2e-4 & log2(fold-change) > 1) in orange and bait in pink. (B) Schematic of Pho7 protein domain organization with DBD in orange and optimal 14-3-3 binding motifs as black bars. (C) UpSet plot indicating number of TF IPs with significant Rad24 and Rad25 enrichments (IP vs untagged; adj. p-value < 0.01 & log2(fold-change) > 1). Colored by optimal 14-3-3 protein binding motif found in TF (pink: TRUE, orange: FALSE). (D) and (E) Interaction networks for the 150 and 500 mM NaCl IP-MS screens, respectively, with baits as orange squares. Physical interactions of TFs not annotated on PomBase^14^ (February 2024) in pink. Similarity score cutoff 6. Interactive version available in TFexplorer (show in [TFexplorer](https://data.fmi.ch/TFexplorer/pombe.html#Network)). See also Figure S6.

At least two other TFs, Prz1, a calcineurin signaling pathway TF, and Ste11, a key TF in sexual differentiation, are negatively regulated by direct 14-3-3 protein interactions^91,92^, suggesting a broader role for them in TF regulation. To explore this, we compared all IPs to an untagged control, focusing on identifying interactors common to many TFs (Figure S6A). This revealed 24 TFs co-purifying with one or both *S. pombe* 14-3-3 proteins (adjusted p-value < 0.01 & log2(fold-change) > 1) (Figure 6C). Though largely of unknown phosphorylation status, ten of these TFs have motifs matching the optimal 14-3-3 binding site (R-x-x-[S/T]-x-P), as identified by ScanProsite^93^ (Figure 6C, Figure S6B), with many more potentially interacting through derivatives of this motif^94^.

These findings highlight abundant interactions between TFs and 14-3-3 proteins, offering novel mechanistic explanations for the Pho7-mediated phosphate starvation stress response and suggesting a widespread and possibly conserved regulatory mechanism for TFs.

### Interactively explorable networks to display putative prey-prey interactions

To complement TF-specific interactions with a global analysis of protein interactions, we extracted bait-agnostic information from our dataset and visualized the results of our IP-MS screens as a network representation of putative PPIs (Figure 6D, Figure 6E). By computing pairwise similarities among all enriched proteins for both low and high salt IP-MS screens (Figure S6C), we gained insights into proteins exhibiting strong correlations across one or multiple IPs. Because this approach is bait-agnostic and our dataset is large, it allows us to detect protein correlations beyond the TFs, revealing additional prey-prey connections. For example, subunits of the same complex that are consistently co-purified together contribute to these connections. While edges between vertices may indicate potential direct physical interactions, they can also represent indirect interactions resulting from shared interactions with other proteins. In these networks, we marked all TF interactions that are not annotated as physical interactions on PomBase^14^ as of February 2024 (Figure 6D, Figure 6E). This includes unannotated yet known TF interactions, such as Scr1’s interaction with the transcription coregulator complex Ssn6/Tup^95^, or associations between TFs and all identified complex members, where the interaction interface likely involves only a few subunits. Due to space constraints, we have set a high threshold for displayed interactions, however, an interactive version of these networks is available in the TFexplorer (show in [TFexplorer](https://data.fmi.ch/TFexplorer/pombe.html#Network)). These networks allow the community to explore interactions with TFs and could reveal variations in cofactor subunit compositions or potentially even lead to the discovery of new cofactor complexes.

### The Nattou complex represses two transmembrane transporter genes and is linked to perinuclear gene localization

Our study uncovered an interaction between two binuclear zinc cluster TFs, Ntu1 and Ntu2, that we propose to form a stable heterodimeric complex (Figure 7A). Ntu1 and Ntu2 feature characteristic coiled-coil structures adjacent to their DBDs, presumably facilitating protein dimerization, and contain fungal-specific MHR domains that are proposed to regulate TF activity^49^ (Figure 7B). Their heterodimerization is further supported by an AlphaFold^96–98^ prediction indicating interaction surfaces at the dimerization domain and between their MHRs (Figure 7B, Figure S7A). Additionally, we observed highly similar gene expression profiles between the individual KO strains when we assessed their role in transcriptional regulation using mRNA-seq (Figure 7C). Particularly, the expression of two predicted transmembrane transporter genes^14^, *tna1^+^* and SPCC576.17c, was strongly upregulated in the absence of either Ntu1 or Ntu2, indicating highly specific and mutually dependent gene repressive activities. Consistent with this observation, our ChIP-seq data revealed that both TFs bind to the promoters of *tna1^+^* and SPCC576.17c, with a complete loss of binding for each TF in the respective KO strain (Figure 7D). These results indicate that Ntu1 and Ntu2 jointly repress *tna1^+^* and SPCC576.17c transcription.

**Figure 7.**
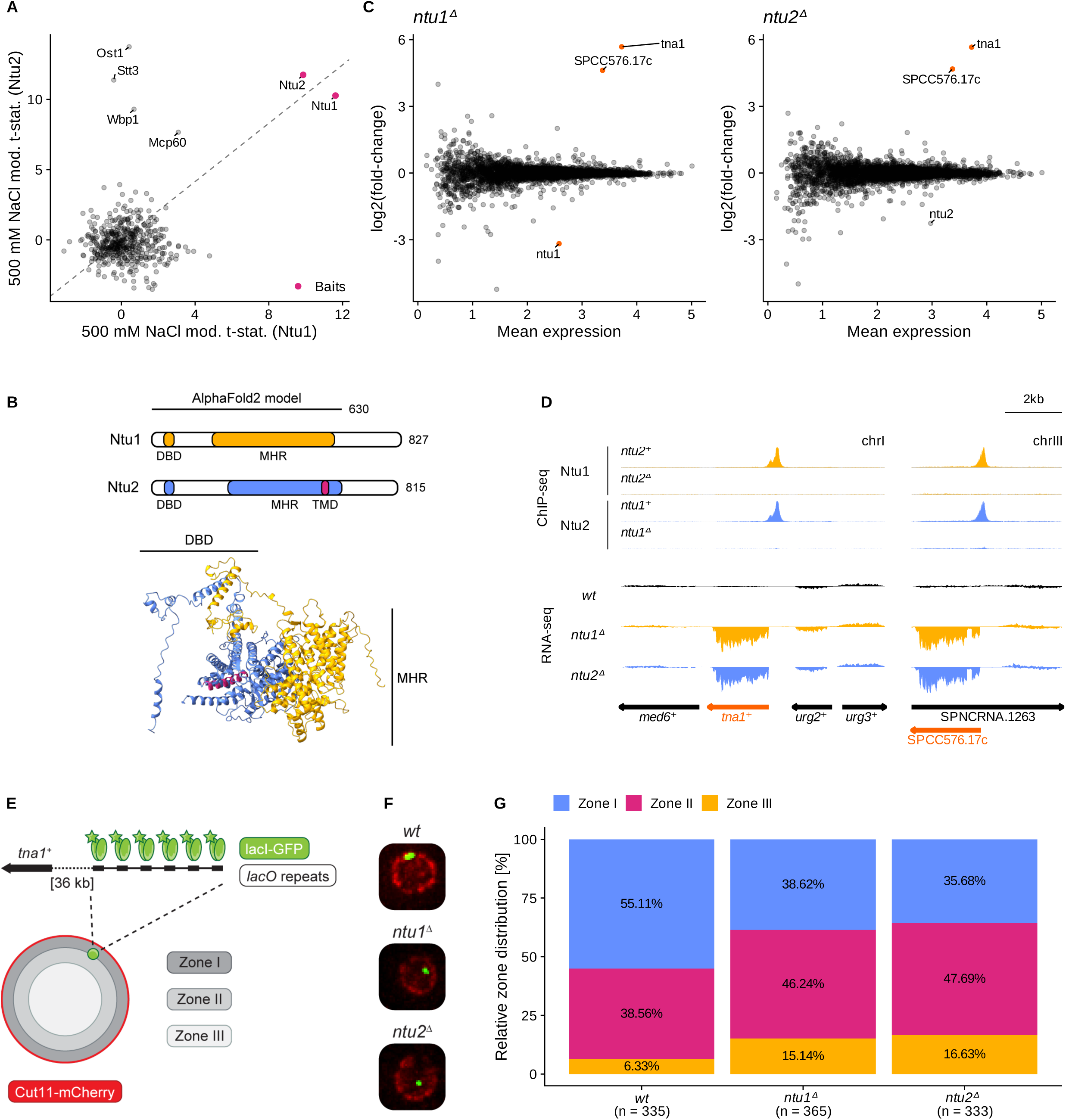
The Nattou complex represses two transmembrane transporter genes and is linked to perinuclear gene localization. (A) Scatter plot correlating moderated t-statistics of proteins in Ntu1 and Ntu2 500 mM NaCl IP-MS experiments, with baits in pink. (B) Schematic of Ntu1 and Ntu2 protein domain organization (top). Cartoon model of the Ntu1/Ntu2 interaction as predicted by AlphaFold^96–98^ including the first 630 amino acid residues with Ntu1 in yellow, Ntu2 in blue, and Ntu2’s predicted TMD in pink (bottom). (C) Differential expression analysis of *ntu1^Δ^* or *ntu2^Δ^* cells compared to wild type. Genes with absolute log2(fold-change) > 3 & adj. p-value < 0.001 in orange. (D) Genome browser views at the *tna1^+^* (chrI) and SPCC576.17c (chrIII) loci. Top: Relative ChIP-seq fragment densities for Ntu1 (yellow) and Ntu2 (blue) in wild-type and backgrounds. Input overlay in grey. Scale: CPM per bp, subset to the loci of interest and scaled by dividing by the largest CPM value, separately for each of the two displayed loci and for each anti-FLAG ChIP. Bottom: Relative mRNA-seq fragment densities (see methods) of *wt*, *ntu1^Δ^*, and *ntu2^Δ^* cells. (E) Schematic representation of microscopy setup to monitor *tna1^+^* locus (marked by lacI-GFP binding *lacO* array 36 kb downstream) relative to the nuclear envelope (marked by Cut11-mCherry). The nucleus is divided into three concentric zones of equal surface area (Zones I-III). (F) Single images of maximum intensity Z-stack projections of *wt*, *ntu1^Δ^*, and *ntu2^Δ^* cells. (G) Quantification of the relative zone distribution of lacI-GFP in *wt*, *ntu1^Δ^*, and *ntu2^Δ^* cells. n indicates the number of cells counted in two independent experiments. See also Figure S7.

A notable feature of Ntu2 is a predicted transmembrane domain (TMD) at its C-terminus^99^ (Figure 7B). We hypothesized that the TMD could tether Ntu2 to the inner nuclear membrane, anchoring its target locus at the nuclear periphery. To test this, we used a strain harboring a *lacO* array close to the *tna1^+^* locus and monitored its positioning within the nucleus relative to the nuclear envelope (Cut11-mCherry) in both wild-type and Nattou KO cells by expressing lacI-GFP (Figure 7E). Dividing the nucleus into three concentric zones of equal surface area, with Zone III representing the nuclear center, we found the *lacO* array predominantly located near the nuclear periphery (Zone I) in wild-type cells (Figure 7F, Figure 7G), consistent with our hypothesis. In contrast, we observed a shift of the locus toward the nuclear center in Ntu1 or Ntu2 KO cells (Figure 7F), which was quantifiable across hundreds of cells (Figure 7G, Figure S7B), indicating Nattou-dependent subnuclear localization. Specifically, the percentage of cells with the *lacO* array at the nuclear periphery (Zone I) decreased from 55% in the wild type to 39% and 36% in *ntu1^Δ^*and *ntu2^Δ^* cells, respectively. Conversely, the proportion in the nuclear center (Zone III) increased from 6% in the wild type to 15% and 17% in the respective KOs.

These results suggest that Ntu1 and Ntu2 form a stable heterodimeric complex and codependently repress two predicted transmembrane transporter genes. Under optimal conditions, one of these genes predominantly resides at the nuclear periphery in a Nattou-dependent manner, potentially mediated by the TMD of Ntu2, implicating that subnuclear localization may be important for the repressive activity of Nattou.

## DISCUSSION

In this study, we addressed gaps in resources and knowledge by creating a strain collection of 89 predicted *S. pombe* TF genes, each tagged endogenously with a FLAG epitope. This collection is available through the National Bio-Resource Project (NBRP) – Yeast, Japan [FY49691-FY49780]. Using this library, we constructed a comprehensive atlas of TF interactions with proteins and chromatin in vegetatively growing cells under optimal conditions. We identified putative protein interactors for approximately half of the investigated TFs, with over a quarter potentially forming stable complexes. Additionally, we discovered potential DNA binding sites for most TFs, covering 2,027 unique genomic regions and revealing motifs for 38 TFs. Our findings offer valuable insights into gene regulation in *S. pombe* and highlight biological phenomena with broader relevance, such as HOT regions and TF regulation by 14-3-3 proteins. To facilitate independent evaluation and further research, we have made all data, metadata, and code accessible, ensuring full reproducibility of our analyses. All figures and numbers can be reproduced using the reports available on GitHub (see [Reports](https://fmicompbio.github.io/Spombe_TFome/#links-to-code-for-generating-figures)). Furthermore, we developed TFexplorer, an interactive web application that allows users to explore the datasets without computational expertise, intended to complement ongoing studies and inspire new research directions.

### Genomic areas with a high density of TF binding

In our comparative analysis of all ChIP-seq experiments, we identified genomic regions occupied by many TFs, exhibiting high accessibility and distinct enrichments similar to HOT regions reported in other organisms^25–27^. We propose that these regions in *S. pombe* also be referred to as HOT regions. Notably, a core set of *S. pombe* TFs was found at most HOT regions (Figure 2C), indicating a unique regulatory environment. Although motif-driven DNA occupancy alone cannot fully explain these regions, one-third contain the DNA-binding motif for Atf1/Pcr1 (show in [Report](https://fmicompbio.github.io/Spombe_TFome/ms_figure_3_chip.html#Facts_and_numbers_-_ChIP-seq)), two TFs frequently enriched at HOT regions. Intriguingly, Atf1 and Pcr1 co-purify with core histone proteins under stringent conditions (Figure 5C), hinting at pioneering TF activity which may be required for establishing these highly accessible regions. Thus, Atf1 and Pcr1 could serve as starting points for further investigations into HOT region characteristics. Additionally, we found that Rst2, another TF enriched at these regions, interacts with the multimodal co-activator complexes SAGA and NuA4 (Figure 5D), suggesting that examining the role of chromatin modifiers could be another avenue for understanding HOT region formation and function.

### Functional interplay between TFs and chromatin regulators

The extent to which TFs specify the activity of chromatin-modifying complexes via direct physical interactions is not well understood. Our study identified limited, rather weak interactions between TFs and known cofactors, suggesting that direct recruitment of chromatin-modifying complexes in *S. pombe* might be uncommon or transient, without persistent TF-cofactor binding. One notable exception is the interaction between three TFs and the acetyltransferase complexes SAGA and NuA4 during low stringency purifications (Figure 5D). Given the ubiquitous role of SAGA and NuA4 in gene activation, broader TF interactions might be expected. Instead, our results align with a study by Göös *et al.*^7^, which found that only a small subset of human TFs specifically interact with SAGA and NuA4.

We also identified a stable interaction between the TF Moc3 and the allantoicase Dal2 (Figure 5C), an enzyme typically involved in purine metabolism, suggesting a non-canonical role for Dal2 in transcriptional regulation. A recent preprint proposed a moonlighting function for an allantoicase protein in sexual reproduction in the malaria parasite *Plasmodium berghei*^100^. Future investigations could explore the intriguing possibility of a role for Dal2’s carbon-nitrogen bond hydrolase activity on chromatin, potentially guided by a sequence-specific TF.

Together, our results suggest a potentially new chromatin regulatory activity and indicate that TFs in *S. pombe* might generally interact with cofactors transiently. Alternatively, cofactors may not physically interact with most TFs, instead engaging at specific loci as a consequence of TF binding.

### Complex fission yeast TF regulatory networks

Our large-scale ChIP-seq analysis revealed that approximately 27% of *S. pombe* protein-coding genes are bound by TFs under optimal growth conditions. Bound genes exhibited slightly higher expression levels, yet non-bound genes were not transcriptionally silent (Figure S2C). This suggests that certain genes may be expressed independently of specific TFs or may become TF-bound only in response to specific stimuli. Notably, several *S. pombe* TFs, such as Ste11, Mbx2, Cbf12, Mei4, and Ace2, are known to autoregulate when active^37,57,101,102^. Our analysis identified eleven additional TFs potentially autoregulating under optimal conditions in vegetatively growing cells (Figure 3), highlighting autoregulation as a prevalent mechanism in *S. pombe*. Interestingly, TF binding was observed up- and downstream of TSSs, as well as just downstream of the transcription end site, indicating regulatory roles beyond the promoter and potential position-dependency. We expect these findings to foster future research into TF binding preferences and position-specific functional properties.

### Possibly perinuclear gene repression by the Nattou complex

We uncovered a stable interaction between two previously uncharacterized binuclear zinc cluster TFs, Ntu1 and Ntu2, which we propose form a repressive heterodimeric complex that we refer to as Nattou (Figure 7). Their similar genome-wide DNA binding patterns and knockout mRNA expression profiles support this hypothesis. One of the Nattou-repressed genes, *tna1^+^*, preferentially localizes near the nuclear periphery in a Nattou-dependent manner, possibly due to Ntu2’s predicted TMD. This localization is interesting, as the nuclear periphery has been associated with reduced transcription and gene silencing^103^. However, Ntu1/Ntu2-mediated repression is restricted to *tna1^+^*(Figure S7C, Figure S7D), which suggests that transcriptional regulation at this locus may not exclusively depend on its subnuclear position. Further research will be needed to validate this hypothesis. We attempted to generate a separation-of-function *ntu2* allele by deleting the Ntu2 TMD to test its role in DNA binding independently of a potential nuclear envelope association. However, this deletion abolished binding to the *tna1^+^* promoter (Figure S7E), similar to the KO phenotype. It is possible that Nattou binding or assembly require attachment to the nuclear envelope. Alternatively, the loss of chromatin binding might result from protein instability, structural changes in the DBD, or impaired nuclear import. Despite these challenges, exploring the separation-of-function approach further could provide valuable insights into the causal relationship between gene regulation and subnuclear localization.

### Limitations of this study

One limitation of this study concerns the functionality of the tagged TFs. Given the largely unknown functions of many TFs, ruling out a potential impact of tagging on TF activity with certainty is challenging. Examples where we observed TF impairment were limited: while the interaction between Adn2 and Adn3 was maintained under high salt conditions with tagged Adn2, it was not observed with tagged Adn3, and four C-terminally tagged TFs showed compromised cellular growth, prompting us to re-tag them at the N-terminus. We recommend that researchers carefully evaluate the functionality of tagged TFs prior to studying them in detail. Furthermore, our TF interactome screen relies on a manually curated list of predicted TFs (PomBase^14^, March 2024), which may include proteins not directly involved in transcription regulation or that bind RNA instead of DNA. For instance, SPAC25B8.11 co-purified with an RNA-binding protein even under high stringency. Additionally, our focus on high-confidence interactions might exclude more frequent ones, like interactions with 14-3-3 proteins, and we advise comparing results with untagged controls to identify these. Finally, our study may not capture transient interactions or those dependent on specific growth conditions or cell cycle stages, which may require alternative experimental approaches.

## Supporting information

Key Resources Table

## ACKNOWLEDGEMENTS

We thank the members of the Bühler lab for their discussions and for testing the beta version of the TFexplorer. Special thanks go to Yukiko Shimada and Nathalie Laschet for their technical support. We thank Martina Donati for the design and development of the TFexplorer web application and her support. We also thank all members of the FMI genomics and mass spectrometry technical platforms for their constant support, and the FMI IT team for hosting TFexplorer. We are grateful to Nadine Vastenhouw, Helge Grosshans, Dirk Schübeler, Fabio Mohn, Jakob Schnabl, Josip Ahel, and Lisa Baumgartner for their comments on the manuscript. This work was supported by the Novartis Research Foundation. S.B. is supported by the Heisenberg Programm of the German Research Foundation (DFG: project ID 464293512). V.N.S.S. was supported by the MSCA ITN Cell2Cell fellowship (project ID 860675).

## Author contributions

Conceptualization, M. Skribbe, C.S., M.B.S., M.B.; Data curation, M. Skribbe, C.S., M.B.S., V.N.S.S., J.S.; Formal Analysis, M. Skribbe, C.S., M.B.S., V.I., D.H., J.S., S.A.S.; Funding acquisition, M.B.; Investigation, M. Skribbe, C.S., M.B.S., V.N.S.S., V.I., D.H., E.P.F.M.; Methodology, M. Skribbe, C.S., M.B.S.; Project administration, M. Skribbe, M.B.; Resources, S.B., M.B.; Software, C.S., M.B.S., M. Schwaiger; Supervision, S.B., M.B.; Validation, M. Skribbe, C.S., M.B.S., V.N.S.S.; Visualization, M. Skribbe, C.S., M.B.S., M. Schwaiger; Writing – original draft, M. Skribbe; Writing – review & editing, M. Skribbe, C.S., M.B.S., M. Schwaiger, V.N.S.S., S.B., J.S., S.A.S., M.B.

## Declaration of generative AI and AI-assisted technologies in the writing process

During the preparation of this work the authors used ChatGPT in order to improve the readability and language of the manuscript. After using this tool, the authors reviewed and edited the content as needed and take full responsibility for the content of the published article.

## METHODS

### Resource availability

#### Lead contact

Further information and requests for resources and reagents should be directed to and will be fulfilled by the lead contact, Marc Bühler.

#### Materials availability

The yeast strain library generated in this study has been deposited to National Bio-Resource Project (NBRP) – Yeast, Japan [FY49691-FY49780].

#### Data and code availability

Sequencing data from ChIP-seq and mRNA-seq have been deposited at GEO (accession numbers GSE274238 and GSE274240) and are publicly available as of the date of publication. Accession numbers are listed in the key resources table. The mass spectrometry proteomics data have been deposited to the ProteomeXchange Consortium via the PRIDE^107^ partner repository with the dataset identifier PXD054070, including einprot^108^ reports with the processing of MaxQuant outputs. Microscopy data reported in this paper will be shared by the lead contact upon request. This paper analyzes existing, publicly available data. Their accession numbers are listed in the key resources table.

All original code has been deposited on Zenodo (https://zenodo.org/records/13270429) and is publicly available as of the date of publication. In addition to the code this archive includes any additional information required to reanalyze the data reported in this paper with individual reports to reproduce all figures and numbers reported in this study and detailed explanations of the code. Also available on GitHub at https://fmicompbio.github.io/Spombe_TFome/. DOI is listed in the key resources table.

Any additional information required to reanalyze the data reported in this paper is available from the lead contact upon request.

### Experimental model and study participant details

#### S. pombe strains and growth conditions

All experiments in this study were conducted using haploid cells of the fission yeast *S. pombe*. Strains were generated using the standard genetic methods of DNA transformation or standard mating^109,110^. Cells used for all experimental procedures were grown in rich medium (YES) liquid culture at 30°C until mid-log phase. Table S1 contains all information on the strains used in this study.

### Method details

#### Selection of TFs for investigation

Based on the curated list of 93 TFs from the fission yeast model organism database PomBase^14^ (as of March 2024), we selected 89 TFs for investigation (go to [File](https://github.com/fmicompbio/Spombe_TFome/raw/main/data/TF_DBD_annotation.txt)). This excludes four TFs (mat1-Mi, mat1-Mc, mat2-Pi, mat3-Mc), as they are associated with the fission yeast mating types^16^ *h^+^* and *h^-^* and our TF interaction screen is conducted in haploid, homothallic *h^-^*strains.

#### Yeast strain generation and culturing

All *S. pombe* strains used in this study are listed in Table S1. Library strains were generated using the PCR-based method^109^ and each strain expresses a fusion protein of a putative TF tagged with a 3xFLAG tag separated by a 6xGlycine linker. All but four library strains were tagged at the C-terminus and carry a *kanMX* selection marker. Cbf11, Mbx2, Pcr1, and Php2 were tagged at the N-terminus using the *ura4*-based gene disruption and replacement system^111^ following unsuccessful genome editing for a C-terminal tag. Successful genome editing of all strains was verified by genotyping PCR and, for tagged strains, by western blot analysis. In addition, we validated all tagged strains used for ChIP-seq by extracting all reads from ChIP input samples that did not map to the *S. pombe* (wild-type) reference genome and aligned them to reference sequences corresponding to all tagged TFs (see methods Genomics data analysis, Alignment). In gene deletion strains the open reading frame was replaced either by a *kanMX* or *natMX* resistance marker. PCR templates for homologous recombination were generated from pFA6a plasmids and include 80 bp sequence homology to the edited locus on both sides (see Table S2 for primers and Data S1 for plasmids). All homology templates were amplified using the NEBNext® High-Fidelity 2X PCR Master Mix (NEB). Strains used in the microscopy assays were generated using the PCR-based method and standard mating^109,110^. Fission yeast strain FY38775^112^ was provided by the National Bio-Resource Project (NBRP) – Yeast, Japan and contains the *lacO*:lacI-GFP reporter at position *chrI*: 1,866,360. All strains were selected on YES plates containing the respective antibiotic. For experiments, cells were cultured in rich medium (YES) liquid culture at 30°C until mid-log phase.

#### Immunoprecipitation

Per replicate, 50 ml of yeast culture was grown to mid-log phase (OD_600_ approximately 1.0) and harvested at 1,200 rcf at 4°C for 4 min. Cells were washed twice with 5 ml cold TBS (50 mM Tris-HCl pH 7.5, 150 mM NaCl) before resuspension in 500 *μ*l cold TBS and transfer to a 2 ml screw-cap tube. Cells were harvested at 3,300 rcf at 4°C for 30 s. The supernatant was removed, and cell pellets were flash-frozen in liquid nitrogen (storage at −80°C until further processing). Initially, IPs were performed using a tube-based protocol (for 30 TFs at 150 mM NaCl) before we adapted the protocol to a 96-well plate format for all subsequent IP experiments to increase throughput, in the following distinguished by “plate” and “tube” annotations.

Frozen cell pellets were thawed on ice and cells were disrupted in the screw-cap tubes in 200 *μ*l (plate)/ 400 *μ*l (tube) Lysis Buffer (150 mM NaCl (low salt) or 500 mM NaCl (high salt), 20 mM HEPES pH 7.5, 5 mM MgCl_2_, 1 mM EDTA pH 8.0, 10% Glycerol, 0.25% Triton X-100, 0.5 mM DTT (fresh), 1x HALT Protease Inhibitor Cocktail (Thermo Fisher Scientific)) with Zirconia/Silica beads (0.5 mm, BioSpec) up to the cell suspension meniscus, using a FastPrep-24 5G bead beating grinder (MP Biomedicals) for three rounds of 20 s at 6.5 m/s. In between bead beating rounds, tubes were cooled on ice for 3 min to avoid overheating.

To collect the crude lysate, tube bottoms were punctured with a 26G needle, placed into 5 ml round bottom polystyrene tubes, and centrifuged two times at 196 rcf at 4°C for 1 min. Beads were shaken loose in between rounds. The lysate was cleared in 1.5 ml reaction tubes by centrifuging twice at 16,100 rcf at 4°C for 10 min. After the final clearing step, the cleared lysate was transferred to a 96-well PCR plate (non-skirted) (plate) or a fresh 1.5 ml reaction tube (tube). Plate-based assay note: For parallel processing of 36 IPs (12 samples in triplicates), each set of replicates was lysed separately, and cleared lysates were kept in the covered 96-well plate on ice until all samples were processed.

Per IP, 2 *μ*g (plate)/ 2.5 *μ*g (tube) anti-FLAG M2 antibody (Sigma-Adrich) were pre-coupled to 15 *μ*l (plate)/ 30 *μ*l (tube) Dynabeads Protein G (Thermo Fisher Scientific) in TBS-T (0.02% Tween 20) for 30 min at room temperature and incubated with the cell lysate for 2 h at 4°C, rotating. Plate wells were sealed with lids and not foil.

Beads were separated on ice using the MagnaBot® II Magnetic Separation Device (Promega) (plate) or the DynaMag™-2 Magnet (Thermo Fisher Scientific) (tube). After the beads settled (5-10 min), they were washed with cold buffers. Plate-based assay note: Buffer addition with a multi-channel pipet, 180° plate rotation for beads to travel through the buffer, plate incubation on ice for 3 min away from the magnetic plate, bead collection on the magnetic plate, and careful removal of the buffer with a multi-channel suction tool.

Beads were washed twice with 200 *μ*l (plate)/ 500 *μ*l (tube) Lysis Buffer (without protease inhibitor) and twice with 200 *μ*l (plate)/ 500 *μ*l (tube) Wash Buffer (100 mM NaCl, 20 mM HEPES pH 7.5, 5 mM MgCl_2_, 1 mM EDTA pH 8.0, 10% Glycerol, 0.25% Triton X-100). To remove all detergent, beads were resuspended in 200 *μ*l B100nd Buffer (100 mM NaCl, 10 mM Tris-HCl pH 7.5, 2 mM MgCl_2_) and transferred to fresh wells/ tubes. After beads settled (5-10 min), the supernatant was removed and beads were washed once more with 200 *μ*l B100nd Buffer for 5 min (for B100nd washes, plate/ tubes are not removed from the magnet and the buffer is removed with a pipet instead of the suction tool).

Beads were digested for 2-3 h at 22°C (shaking occasionally) with 0.2 *μ*g Lys-C (Wako) (1 *μ*l) in 5 *μ*l digestion buffer (3 M GuaHCl, 20 mM EPPS pH 8.5, 10 mM CAA, 5 mM TCEP). Plate wells were sealed with fresh lids. Samples were diluted with 17 *μ*l 50 mM HEPES pH 8.5 prior to the addition of 0.2 *μ*g Trypsin (Thermo Fisher Scientific) (1 *μ*l) and incubated over-night at 37°C. The next day, the supernatant was transferred to a 96-well plate (skirted) for LC-MS/MS analysis.

IP-MS experiments for each TF and the parental untagged controls were conducted in triplicates. For each batch (11x 150 mM NaCl IP-MS; 4x 500 mM NaCl IP-MS) of IP-MS experiments, a positive control (Atf1-3xFLAG) and negative control (untagged) was included. For the analysis, only one of each control per experimental protocol was included (see Quantification and statistical analysis, Proteomics data analysis).

#### Mass spectrometry

Peptides generated by Lys-C/ Trypsin digestion were acidified with 0.8% TFA (final concentration) and analyzed by LC–MS/MS on either an EASY-nLC 1000 or a Vanquish Neo chromatography system (both Thermo Fisher Scientific) with a two-column setup (*μ*PAC trapping column and 50 cm *μ*PAC column, or a 0.3 x 5 mm Pepmap C18 trapping column and an EASY-Spray Pepmap Neo 2 *μ*m C18 75 *μ*m x 150 mm column heated to 45°C) mounted on an EASY-Spray™ source connected to an Orbitrap Fusion Lumos mass spectrometer (all Thermo Fisher Scientific, or formerly Pharmafluidics). The peptides were loaded onto the trapping column a in 0.1% formic acid and 2% acetonitrile in H_2_O, then separated at room temperature (*μ*PAC) or 45°C (Pepmap Neo) over an 80 min gradient with a mobile phase buffer system consisting of Buffer A: 0.1% formic acid; Buffer B: 0.1% formic acid in 80% acetonitrile in a linear gradient of 2–7% Buffer B in 3 min followed by a linear increase from 7-20% in 45 min, 20–30% in 15 min, 30–36% in 8 min, 36-45% in 2 min, and the column was finally washed for 7 min at 100% Buffer B.

MS1 survey scans were performed every 3 s using a 120k resolution (at 200 m/z) in the Orbitrap from 375-1575 m/z in profile mode, with a maximum injection time of 50 ms. Precursors for MS2 were selected using advanced peak determination and monoisotopic peak determination was set to peptides and charge states 2-7 were allowed for fragmentation. A dynamic exclusion of 8 s within +/-10 ppm tolerance was used, and the minimum intensity was set to 1e4. Precursors were selected in the quadrupole within an isolation window of 1.6 m/z for HCD (higher energy collisional dissociation) fragmentation. The normalized collision energy was set to 35%. A maximum injection time of 60 ms was used and the ion trap was scanned in rapid mode. MS2 data were acquired in centroid mode.

The mass spectrometry proteomics data have been deposited to the ProteomeXchange Consortium via the PRIDE^107^ partner repository with the dataset identifier PXD054070.

#### Chromatin immunoprecipitation

Per replicate, 50 ml of yeast culture was grown to mid-log phase (OD_600_ approximately 1.0-1.2) and harvested at 1,200 rcf for 4 min. Cells were resuspended in 5 ml PBS (room temperature) and crosslinked with 1% formaldehyde (Sigma-Aldrich) at room temperature for 15 min, rotating. The crosslinking reaction was quenched with 140 mM Glycine for 5min, rotating. Cells were harvested at 1,200 rcf at 4°C for 4 min and washed twice with 15 ml cold PBS before resuspension in 500 *μ*l cold PBS and transfer to a 2 ml screw-cap tube. Cells were harvested at 3,300 rcf at 4°C for 30 s. The supernatant was removed, and cell pellets were flash-frozen in liquid nitrogen (storage at −80°C until further processing).

Frozen cell pellets were thawed on ice and cells were disrupted in 400 *μ*l Lysis Buffer (50 mM HEPES pH 7.5, 140 mM NaCl, 1 mM EDTA pH 8.0, 1% Triton X-100, 0.1% sodium deoxycholate (fresh), 1x HALT Protease Inhibitor Cocktail (Thermo Fisher Scientific)) with Zirconia/Silica beads (0.5 mm, BioSpec) up to the cell suspension meniscus, using a FastPrep-24 5G bead beating grinder (MP Biomedicals) for three rounds of 1 min at 6.5 m/s. In between bead beating rounds, the tubes were cooled on ice for 3 min to avoid overheating. To collect the crude lysate, tube bottoms were punctured with a 26G needle, placed into 5 ml round bottom polystyrene tubes, and centrifuged two times at 196 rcf at 4°C for 1 min. Beads were shaken loose in between rounds. 1 ml of Lysis Buffer was added to the crude lysate before transferring it to 15 ml Bioruptor® Pico Tubes (diagenode) with sonication beads (diagenode). Samples were sonicated twice for 12 cycles (30 s on/ 30 s off) using the Bioruptor® Pico sonication device (diagenode) with 10-12 min incubation on ice in between sonication rounds to avoid overheating. The lysate was cleared in 1.5 ml reaction tubes by centrifuging twice at 16,100 rcf at 4°C for 5 min and 15 min, respectively. After the final clearing step, the cleared lysate was transferred to a fresh 1.5 ml reaction tube. Of each replicate, 50 *μ*l of lysate was added to 50 *μ*l of Lysis Buffer in a fresh tube, set aside as input control, and kept on ice until the de-crosslinking step.

Per IP, 2.5 *μ*g anti-FLAG M2 antibody (Sigma-Adrich) were pre-coupled to 30 *μ*l Dynabeads Protein G (Thermo Fisher Scientific) in TBS-T (0.02% Tween 20) for 30 min at room temperature and incubated with the cell lysate for 2 h at 4°C, rotating.

Beads were separated at room temperature using the DynaMag™-2 Magnet (Thermo Fisher Scientific) and washed with cold buffers. Beads were washed three times with 1 ml Lysis Buffer (without protease inhibitor), one time with 1 ml Wash Buffer (10 mM Tris-HCL pH 8.0, 250 mM LiCl, 1 mM EDTA pH 8.0, 0.5% NP-40, 0.5% sodium deoxycholate (fresh)), and one time with 1 ml 1x TE (10 mM Tris-HCl pH 8.0, 1 mM EDTA). Samples were eluted in 250 *μ*l 1% TES (1x TE, 1% SDS) in two steps. First, beads were incubated in 125 *μ*l 1% TES at 65°C for 10 min and the supernatant was transferred to a fresh 1.5 ml reaction tube. Second, beads were resuspended in 125 *μ*l 1% TES, separated on the magnetic rack, and the supernatant was combined with the previous eluate. 150 *μ*l of 1% TES were added to the input control before ChIP and input samples were de-crosslinked over-night at 65°C.

The next day, samples were treated first with 40 *μ*g RNase A for 1 h at 37°C and then with 60 *μ*g Proteinase K for 1 h at 55°C. DNA was precipitated with 150 mM NaCl and 1 volume of isopropanol and extracted with 30 *μ*l AMPure XP beads at room temperature for 15 min, rotating. Samples were washed twice with 500 *μ*l 80% Ethanol (fresh) on the magnetic rack for 2 min and eluted in 20 *μ*l and 50 *μ*l 1x TE for ChIP and input samples, respectively.

ChIP-seq experiments for each TF and the parental untagged control were conducted from two individual cultures (in duplicates) and each IP sample has a corresponding input sample.

#### ChIP NGS library preparation and sequencing

ChIP-seq libraries were generated using the NEBnext Ultra II DNA Library Prep kit (NEB), according to the manufacturer’s protocol, using NEBnext Multiplex Oligos for Illumina (UDI) (NEB). 10 ng of DNA was used for input samples and the entire material was used for the IP samples, with the upper limit of 10 ng when DNA was quantifiable with Qubit dsDNA (high sensitivity) reagents (Thermo Fisher Scientific). Library concentration was determined using Qubit dsDNA (high sensitivity) reagents (Thermo Fisher Scientific) and average fragment sizes were measured using the Fragment Analyser System (Agilent). Libraries were sequenced using the NovaSeq 6000 platform (Illumina) with paired-end reads (2x 56 bp).

#### mRNA-sequencing

Total RNA was extracted from three individual cultures (triplicates) using the MasterPure Yeast RNA Purification Kit (Lucigen) according to the manufacturer’s protocol. RNA integrity was assessed using the Tapestation RNA ScreenTape reagents kit (Agilent). RNA concentration was determined using the Nanodrop Spectrophotometer (Thermo Fisher Scientific) or Qubit RNA (broad range) reagents (Thermo Fisher Scientific).

mRNA-seq libraries were generated using the Illumina Stranded mRNA Prep (Illumina) according to the manufacturer’s protocol, using 300 ng of total RNA as input. Library concentration was determined using Qubit dsDNA (high sensitivity) reagents (Thermo Fisher Scientific) and the average fragment size of the final pool was measured using the Fragment Analyser System (Agilent). Libraries were sequenced using the NovaSeq 6000 platform (Illumina) with paired-end reads (2x 56 bp).

#### Microscopy

Two independent microscopy experiments were conducted (duplicates). Liquid cultures of *S. pombe* cells were grown over-night in rich medium (YES) at 30°C to the logarithmic phase (OD_600_ approximately 0.6). Prior to imaging, the cells were attached with lectin (Sigma-Aldrich) to glass bottom dishes with a microwell (MatTek). Cells were imaged on a DeltaVision™ Ultra High-Resolution microscope (Cytiva) with an Olympus 60X/1.42, Plan Apo N, UIS2, 1-U2B933 objective. Z-stacks were obtained at focus intervals of 0.25 *μ*m and images were deconvolved with the inbuilt software softWoRx using default settings. Images were randomized before the analysis to avoid bias. For Z-stack processing, the top and bottom three stacks (out of 22 total stacks) were disregarded, and Maximum Intensity Projection was applied. The FiJi/ImageJ software^113^ was used to measure the distances between the foci and the nuclear periphery marked by Cut11-mCherry. For zone-based position quantification the nucleus was divided into three concentric zones with equal surface area (assuming a circular shape of the nucleus).

#### TFexplorer

The data visualization for the TFexplorer webtool (https://data.fmi.ch/TFexplorer/) was done using the LinkedCharts^114^ and IGV.js^115^ libraries in JavaScript.

#### AlphaFold interaction prediction

Structural predictions for the Ntu1/Ntu2 complex were obtained with AlphaFold-Multimer^98^. Jobs were run through GUIFold^116^ which employs a modified pipeline based on AlphaFold2^96,97^ version 2.3.1. Feature generation and prediction was according to the standard Alphafold2-Multimer protocol. Five structural models were predicted using Ntu1 and Ntu2 protein sequences as inputs at a 1:1 stoichiometry. The model with the lowest predicted aligned error (PAE) value for the complete prediction was used for visualization in ChimeraX^117^ version 1.6.1 and colored by protein identity or pLDDT confidence measure.

### Quantification and statistical analysis

#### Proteomics data analysis

##### Identification and quantification

The IP-MS raw files were processed as a single batch with MaxQuant^118^ version 2.2.0.0, with the exception of the Dal2 pulldown experiments which was processed separately, together with a corresponding set of untagged controls. The peptide identification was performed using a fasta database obtained from ENSEMBL (https://ftp.ebi.ac.uk/ensemblgenomes/pub/release-55/fungi/fasta/schizosaccharomyces_pombe/pep/Schizosaccharomyces_pombe.ASM294v2.pep.all.fa.gz) and the built-in MaxQuant contaminants database. MaxQuant was run with mostly default settings, with LFQ (LFQ min. ratio count 2, fastLFQ: True) and iBAQ enabled.

##### Processing

The proteinGroups.txt file produced by MaxQuant^118^ was further processed with einprot^108^ version 0.9.3. Samples generated under low salt and high salt conditions were processed separately. Fully reproducible reports detailing the complete einprot analysis are provided on PRIDE (PXD054070). In summary, potential contaminants, reverse hits, proteins only identified by site, and proteins identified by less than two peptides or with a score below 10 were filtered out. The LFQ intensities were log2- transformed and missing values were imputed using a modified version of the MinProb algorithm implemented in the imputeLCMD R package^119^ version 2.1, sampling values to impute from a normal distribution with parameters derived from all observed values, rather than separately for each sample. Principal component analysis (see Figure S2A) was performed on the imputed log-transformed intensities using the scater^120^ R package version 1.26.1.

##### Statistical testing

The *limma* R package^20^ version 3.54.2 was used to fit a linear model for each group, comparing the imputed log2-transformed intensities from the samples in the group to those in a broad “complement” group. This complement group contained all samples that were generated under the same salt conditions and with the same protocol (tube- or plate-based, see methods Immunoprecipitation) as the group of interest. This approach was selected over, for example, comparing to the untagged controls only, in order to focus the attention on specific interactors rather than non-specific ones, which may be pulled down by many baits. For each comparison, at least two observed (non-imputed) values were required in order to report a test result for a protein, and each sample was assigned a weight inversely proportional to the total number of experiments for the corresponding bait. The adjusted p-values and moderated t-statistics from *limma* were used for most visualizations and determination of interactors. In addition, two other types of comparisons were performed. In order to detect a broader set of interactors (not just specific ones), each group was compared to only the untagged controls obtained with the same protocol. Finally, for increased comparability between low and high salt log2(fold-changes), each low salt group was compared to a complement group made up of the groups that were also studied under high salt conditions (high salt complement) (regardless of the type of protocol used).

##### Generation of interaction networks

The moderated t-statistics obtained from *limma*^20^ were used as the basis for generating protein networks, in which proteins were connected if they showed similar enrichment patterns across the whole set of pulldown experiments. More precisely, the moderated t-statistic profiles were first truncated by setting all values below 0 as well as those corresponding to an adjusted p-value above 0.1 to 0. Next, a similarity score was defined for each pair of proteins. Letting *t̄_ij_* denote the truncated t-statistic for protein *i* in comparison *j*, the similarity between proteins *i* and *k*was defined by

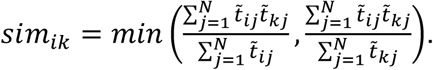

The resulting symmetric matrix of similarity scores (here after thresholding at a suitable similarity value, in the figures in the manuscript set to 6.0) can be interpreted as an adjacency matrix, from which a network can be constructed containing all proteins with at least one remaining edge (for a graphical explanation see (Figure S6C)). The edges in the displayed networks are weighted by the respective similarity scores.

#### Genomics data analysis

##### Alignment

Paired-end reads from ChIP-seq were aligned to the reference genome obtained from ENSEMBL (https://ftp.ebi.ac.uk/ensemblgenomes/pub/release-55/fungi/fasta/schizosaccharomyces_pombe/dna/Schizosaccharomyces_pombe.ASM294v2.dna.toplevel.fa.gz) using QuasR^121^ version 1.38.0 with default parameters, resulting in read pairs with more than a single genomic hit to be discarded. For validation of tagged yeast strains from this study, read pairs that failed to align to the reference genome were further aligned to a custom fasta file (go to File) containing the TF sequences including the added tags. To account for differences in sequencing depth, the counts were normalized by dividing them through the total number of read pairs that mapped to each tagged TF sequence and then visualized as a heatmap (show in Report). BigWig files were generated using QuasR (qExportWig using default parameters and binsize=1, scaling=1e6). Public ATAC-seq data^104^ was first processed by cutadapt^122^ version 3.7 with parameters -a NCTGTCTCTTATA -A NCTGTCTCTTATA --minimum-length=10 --length=50 --overlap=1, and then aligned in the same way as ChIP-seq data (see key resources table for accession numbers of public data).

##### ChIP-seq quantification

Fragments in genomic regions of interest (such as tiles or peaks) were counted using QuasR^121^ with parameters shift=“halfInsert”, orientation=“any”, and useRead=“first”. Raw fragment counts in regions were normalized to log2 counts per million using *lcpm = log*2(*r* / *N* * 1*e*6 + 1), where *lcpm* is the log2-normalized value for a region and sample, *r* is the raw fragment count for a region and sample, and *N* is the total number of aligned fragments in a sample.

##### Identification of problematic regions

Problematic regions (also referred to as blacklisted regions), including regions with low mappability or copy number variations between experimental strains and the reference genome, were identified by first tiling the genome into sequential, non-overlapping windows of 200 bp and quantifying these tiles in all 162 input samples (2 replicates for each of the 80 tagged TF strains and the parental untagged strain). Tile counts were normalized to *lcpm* values and converted to z values in each sample by subtracting the mean and dividing by the standard deviation over tiles on all chromosomes except the mitochondrial (“MT”). Tiles with z values less than −2.575829 (0.5^th^ percentile of a standard normal distribution) or greater than 2.575829 (99.5^th^ percentile of a standard normal distribution) in at least 81 (50%) of the input samples were defined as problematic (2,387 of 63,155 tiles, 3.78%) and excluded from further analyses.

##### Correction of GC-bias in ChIP-seq

Genomic tiles of 200 bp were quantified and normalized to *lcpm* values for all 324 samples. For each sample, a linear model of the form *lcpm* ∼ *gc* was fitted using R’s lm function, where *gc* is the percent G+C bases in a tile. This model can be assumed to mostly capture signals from background (non-enriched) tiles, which typically comprise the large majority of all genomic tiles. The sample-specific coefficients (“GC-slopes”) obtained from these fits, ranged up to values of approximately 0.1 for samples with a clear GC-bias, correspoc input samples, indicating a 2^10 * 0.1^ = 2-fold increase of signal for a 10% increase of G+C content. The GC-slopes were used to calculate GC-corrected counts for regions (tiles or peaks) using 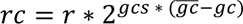, where *rc a*nd *r* are the GC-corrected and raw fragment counts for a region and sample, *gcs* is the sample-specific linear model coefficient (“GC-slope”) and 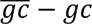 is the difference of a region’s percent G+C from the average percent G+C over all regions. The corrected counts *rc* have a similar magnitude as the raw counts *r* and are then used to calculate log2 counts per million (*lcpm*) as described above. No GC-correction was performed for public data that was not generated in this study.

##### ChIP-seq peak finding and IP enrichment

Peak candidates were identified on the pooled replicates for each TF (IP samples only) using MACS2^123^ version 2.2.7.1 with parameters --gsize 1.21e7 --keep-dup all -- nomodel --shift 0 --qvalue 1e-6. Peak candidates in each sample were set to a width of 500 bp centered on the peak summit identified by MACS2 and then fused across TFs (combining overlapping peaks) to create a common set of 6,102 peak candidates. These peak regions were then quantified in each sample, GC-corrected and normalized to *lcpm* as described above. Enrichments were calculated as *enr* = *lcpm_IP_* − *lcpm_input_* where *lcpm_IP_* and *lcpm_input_* are the *lcpm* values for corresponding IP and input samples, respectively. Peak candidates with enr ≥ *log*_2_ 1.75 in at least two samples (keeping replicates separate) that did not overlap problematic regions were kept for downstream analyses, resulting in a final set of 2,027 peaks. These peaks were classified into three groups: Peaks with *enr* values greater than 1.0 for at most 4 TFs (5%, replicates averaged) were classified as “specific peaks” (1,686 peaks). The remaining peaks were further grouped into “common frequent” (94 peaks) and “common ubiquitous” (247 peaks), using R’s kmeans functions with parameters centers=2, nstart=10 on the *enr* values (replicates averaged). A gene was annotated to be bound by a TF if it had an enriched TF peak (*enr* ≥ *log*_2_ 2.0) overlapping its promoter, defined as a 1 kb bin centered on the annotated transcript start site (TSS annotations of reference genome). Final annotated peaks are provided as comma-separated [File](https://github.com/fmicompbio/Spombe_TFome/raw/main/data/fused_peaks_filtered.csv.gz).

##### Motif finding in ChIP-seq peaks

*De novo* motif identification was performed using STREME^124^ from the MEME motif finding toolbox version 5.5.2. For each of the 62 TFs with at least 2 enriched specific peaks (replicate-averaged *enr* value greater than 1.0), STREME was run with the enriched specific peaks as input (excluding common peaks), all non-enriched specific peaks as control, and parameters objfun=“de”, alph=“dna”, nmotifs=5. The resulting motif candidates were manually reviewed independently by two researchers based on several motif quality criteria (motif occurrence in enriched versus non-enriched peaks and in peaks versus the rest of the genome, motif localization relative to peak mid points, ChIP-seq fragment density near motif hits, and motif similarity to 6-mer words enriched in TF peaks compared to control peaks). The two manual curations were consolidated resulting in a consensus set of 67 motifs (45 unique) identified for 38 TFs (go to [File](https://github.com/fmicompbio/Spombe_TFome/raw/main/data/motifs_annotated.csv)). The universalmotif package^125^ version 1.16.0 was used to calculate motif similarities (compare_motifs function with parameters method = “PCC”, tryRC=TRUE, min.overlap=4, min.mean.ic=0.25, normalise.scores=TRUE) and to scan sequences for motif hits (scan_sequences function with parameters threshold=1e-4, threshold.type=“pvalue”, RC=TRUE).

##### RNA-seq processing

Paired-end reads from RNA-seq were quantified on the transcript level using Salmon^126^ version 1.9.0 with parameters --gcBias --seqBias –numGibbsSamples=50. The reference index was constructed based on reference sequences (cDNA and non-coding RNA sequence) obtained from ENSEMBL (https://ftp.ebi.ac.uk/ensemblgenomes/pub/release-55/fungi/fasta/schizosaccharomyces_pombe/cdna/Schizosaccharomyces_pombe.ASM294v2.cdna.all.fa.gz and https://ftp.ebi.ac.uk/ensemblgenomes/pub/release-55/fungi/fasta/schizosaccharomyces_pombe/ncrna/Schizosaccharomyces_pombe.ASM294v2.ncrna.fa.gz), using the genome sequence

(https://ftp.ebi.ac.uk/ensemblgenomes/pub/release-55/fungi/fasta/schizosaccharomyces_pombe/dna/Schizosaccharomyces_pombe.ASM294v2.dna.toplevel.fa.gz) as a decoy^127^ and a k-mer length of 23. In addition, reads were aligned to the genome using STAR^128^ version 2.7.10b, and the resulting bam files were used as input to DeepTools^129^ bamCoverage version 3.3.1 for generation of bigWig files for visualization. The bigWig files were generated separately for fragments from the positive and negative strand, using a bin width of 1 nt.

##### RNA-seq differential expression analysis

Transcript-level estimated counts from Salmon^126^ were imported into R and summarized to the gene level (only one gene had more than one annotated transcript) using tximeta^130^ version 1.16.1. The fishpond package^131^ version 2.4.1 was used to calculate the inferential relative variance for each gene. Genes with an estimated count of at least 5 in at least two samples, and an average inferential relative variance below 0.178 were retained for differential expression analysis. The DESeq2^132^ package version 1.42.1 was used to create a DESeqDataSet from the SummarizedExperiment object generated by tximeta^130^, and to compare each KO condition to the *wt* control, using default settings except for setting alpha=0.05. MA plots and correlation plots were generated based on the DESeq2 output tables.

##### Coverage plots

Genome coverage plot tracks for the RNA-seq data were generated from the bigWig files created by DeepTools^129^, which were imported into R using the rtracklayer package^133^ version 1.58.0. Coverage of fragments from the positive strand is displayed as positive score values, while coverage of fragments from the negative strand is shown as negative score values. The scores were normalized by the sample-wise library size (the total number of reads assigned to a gene) to generate counts per million (CPM) values, and subset to the regions of interest (on chromosomes I and III, respectively). Relative CPM values, obtained by dividing by the largest observed absolute CPM value, are used for visualization. This normalization is performed separately for the two displayed regions. The coverage tracks of one representative replicate are shown in the figures.

#### Microscopy data analysis

Cell-level measurements of distances of foci from the nuclear periphery as well as the radii of the nuclei were imported into R (go to [File](https://github.com/fmicompbio/Spombe_TFome/raw/main/data/Fig7_lacR_quantification_distance_radius.txt)), and relative distances from the nuclear periphery were calculated. One cell where the estimated distance from the nuclear envelope was marginally larger than the estimated radius was excluded from further analysis. The relative distances were used for the zone-based position classification. Cells where the relative distance to the nuclear periphery was less than 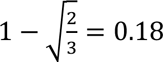 were classified as ‘Zone I’, those with a relative distance larger than 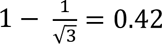 were classified as ‘Zone III’, and the remaining ones as ‘Zone II’. Relative distances were compared between conditions by summarizing the values by the median for each replicate (to avoid pseudoreplication) and fitting a general linear model to the summarized values, with condition as the predictor.

### Additional resources

All data, metadata, and code used in this study is available on GitHub (https://github.com/fmicompbio/Spombe_TFome) including individual reports that allow users to fully reproduce all figures and numbers of this study with detailed explanations of the code.

TFexplorer, an interactive web application that allows users to explore the datasets without computational expertise, is available at https://data.fmi.ch/TFexplorer/.

## SUPPLEMENTAL INFORMATION

Table S1. *S. pombe* strains used in this study, related to STAR Methods

Table S2. Primers used in this study, related to STAR Methods

Data S1. GenBank files of all plasmids used in this study, related to STAR Methods

---

**Figure S1.**
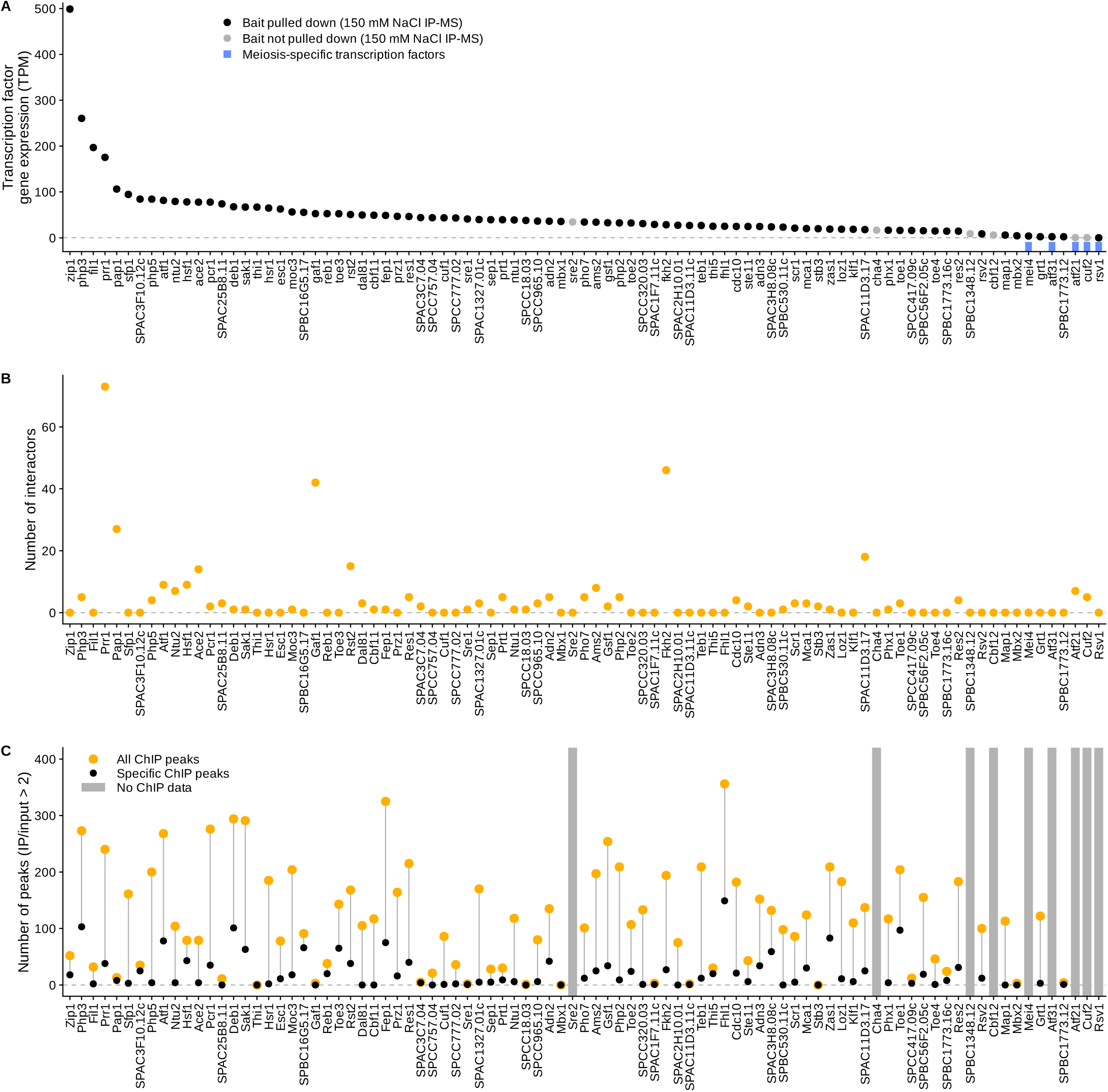
Extended overview of *S. pombe* TF expression and interactome results, related to Figure 1. (A) mRNA expression levels (wild-type) of investigated TFs, ordered by decreasing TPM. Grey dots represent TFs not identified in the low salt IP-MS screen and blue bars indicate meiosis-specific TFs. (B) Number of interactors identified in the 150 mM NaCl IP-MS screen for investigated TFs in the same order as in (A). (C) Number of enriched ChIP peaks (IP/input > 2) for investigated TFs, all peaks in orange and specific peaks (bound by at most four TFs) in black in the same order as in (A). Grey bars indicate TFs not included in the ChIP-seq screen.

**Figure S2.**
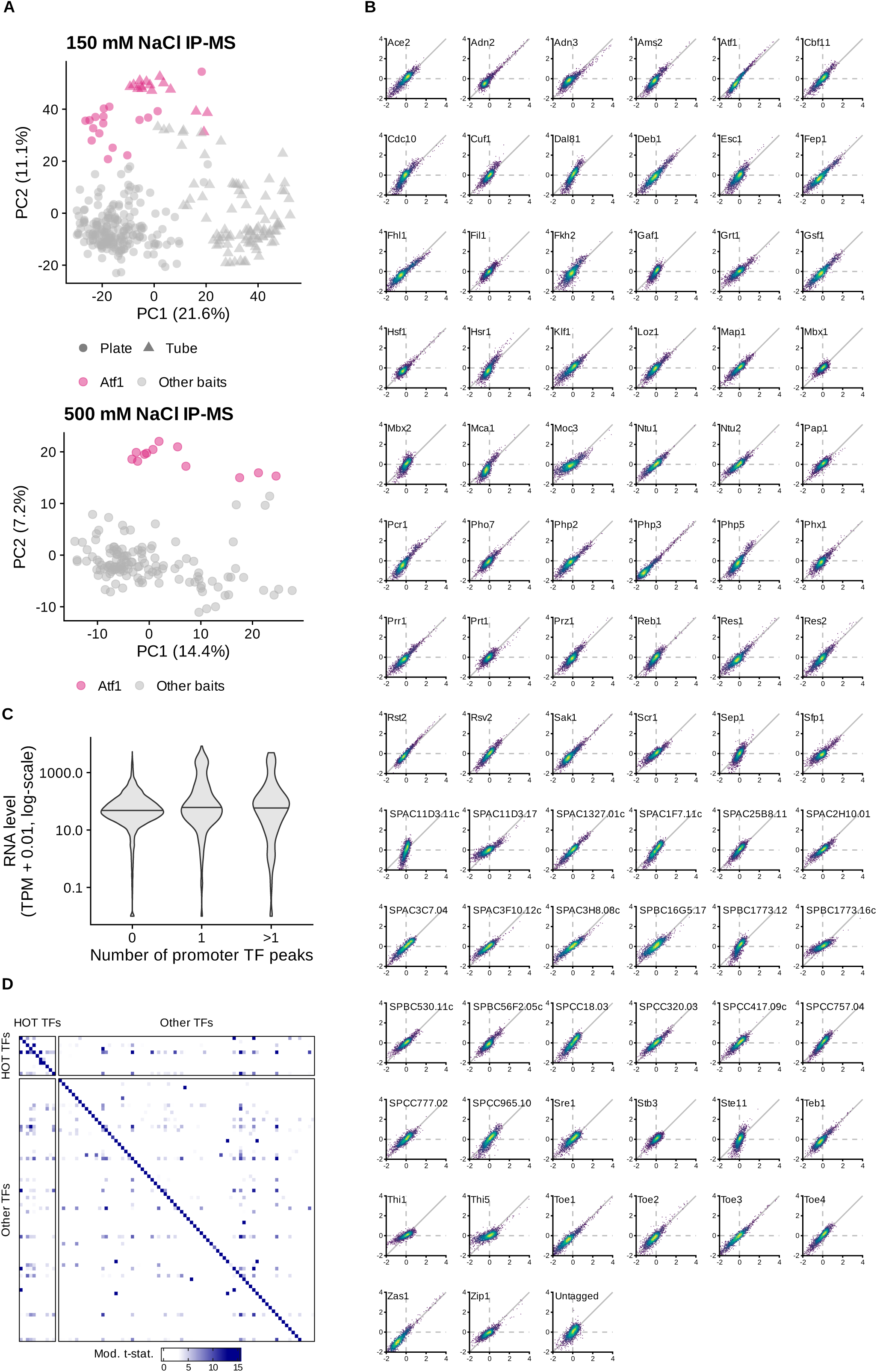
Investigated TFs exhibit abundant protein interactions and diverse genome-wide binding patterns, related to Figure 2. (A) Principal component analysis (PCA) of all IPs at 150 mM NaCl (top) and 500 mM NaCl (bottom). Positive control IPs (Atf1-3xFLAG) in pink. Point shapes indicate experimental protocols (circle = Plate, triangle = Tube). Also see methods. (B) Pairwise correlation of ChIP enrichments (log2(IP/input)) between replicates. (C) Distribution of log-scaled TPM values of protein-coding genes according to the number of TF peaks in their promoter (1 kb bin centered on TSS). (D) Heatmap of moderated t-statistics of the 150 mM NaCl IP-MS screen for all TFs (IP vs untagged control). Heatmap is split by HOT and other TFs, with column order matching row order.

**Figure S3.**
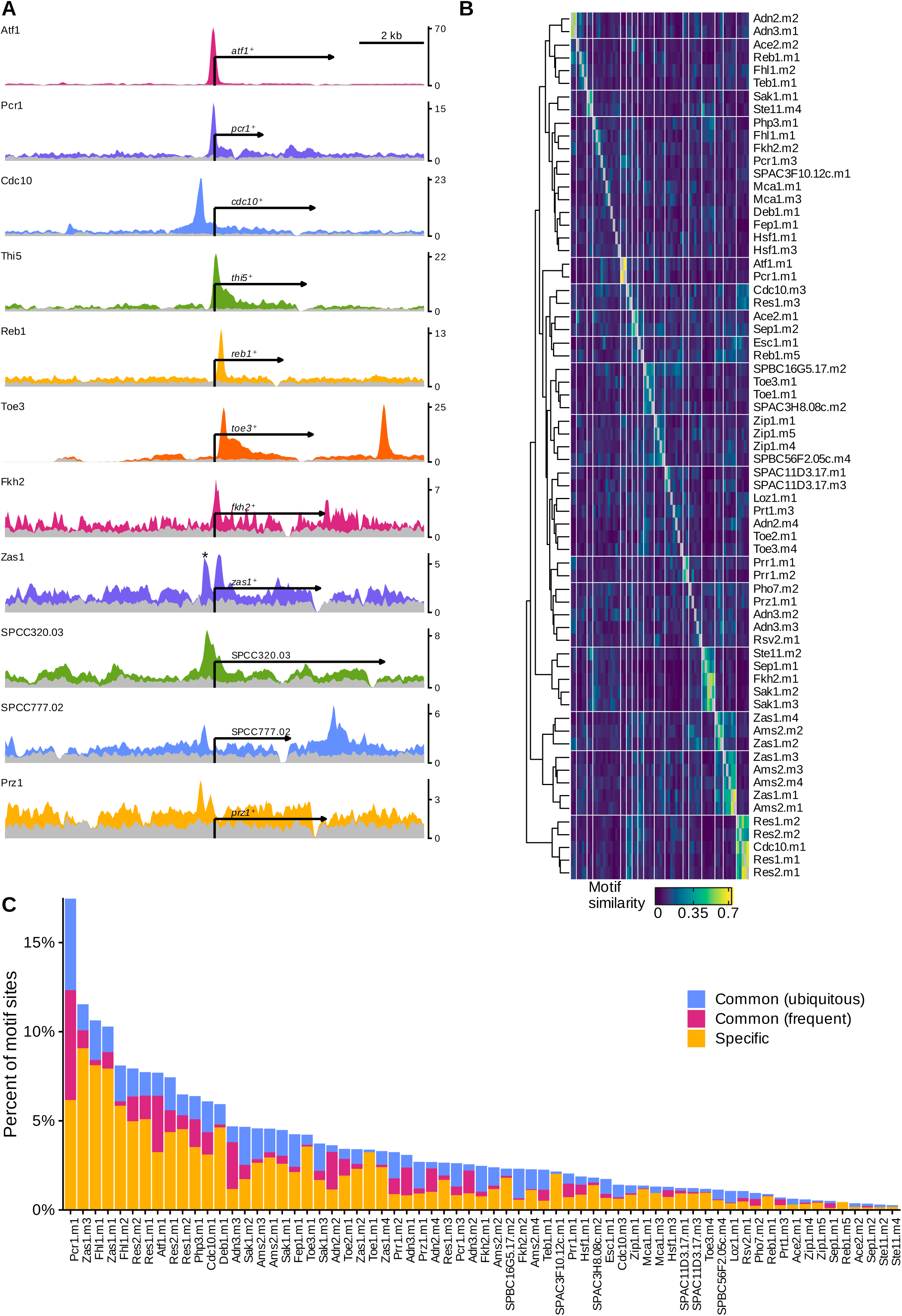
Analysis of TF regulatory networks and *de novo* DNA-binding motif identification, related to Figure 3. (A) Genome browser views of ChIP (colored) and input (grey) fragment densities for eleven TFs around their gene loci. Scale: CPM per bp, smoothed over 101 bp windows using a running mean. Asterisk (*) indicates enrichment over tRNA gene (considered artifact detected in 48 TF ChIPs). All genes are oriented to align at their TSS. Coverage dips at 3’ ends correspond to affinity tag insertion sites. (B) Fully annotated heatmap of pairwise motif similarities (PCC), clustered into 17 groups. (C) Percent of genome-wide motif sites overlapping ChIP enrichments of the corresponding TF, indicated separately for each peak class in blue (common ubiquitous), pink (common frequent), and orange (specific).

**Figure S4.**
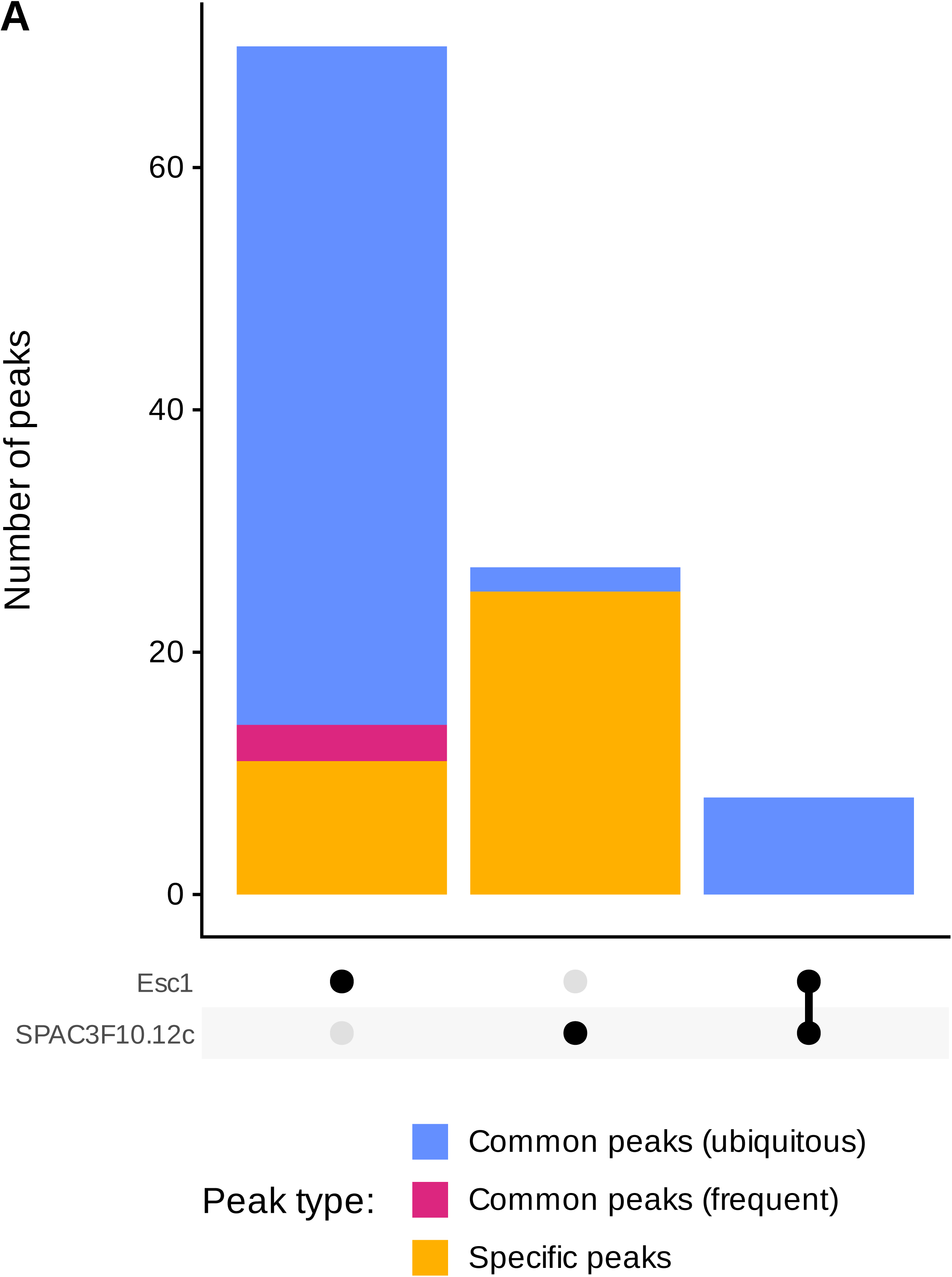
bHLH TF ChIP peaks comparison, related to Figure 4. (A) UpSet plot showing the number of common ubiquitous (blue), common frequent (pink), and specific (orange) ChIP peaks among Esc1 and SPAC3F10.12c.

**Figure S5.**
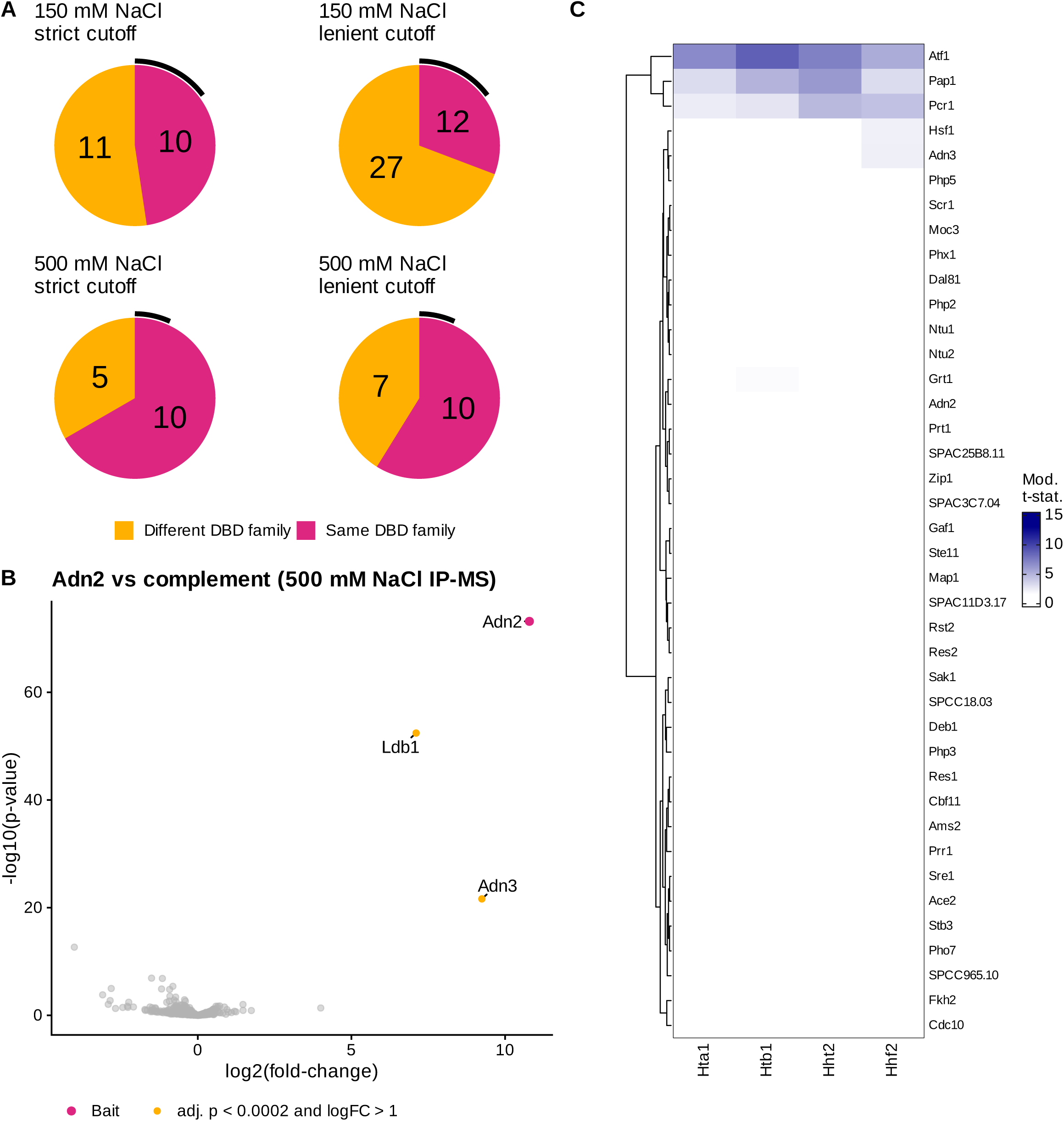
Approximately one in three TFs engage in TF-TF interactions, related to Figure 5. (A) Pie charts of TF-vs-TF interactions detected in the 150 and 500 mM NaCl IP-MS screens qualified using a strict (adj. p-value < 2e-4 & log2(fold-change) > 1) or lenient (adj. p-value < 0.01 & log2(fold-change) > 1) cutoff. Charts are divided into interactions observed within (same) and between (different) DBD families. The black arc indicates the expected fraction of interactions within the same DBD family given the sizes of the DBD families. (B) Volcano plot for 500mM NaCl IP-MS of Adn2 vs complement with significantly enriched proteins (adj. p-value < 2e-4 & log2(fold-change) > 1) in orange and the bait in pink. (C) Heatmap of moderated t-statistics for histone proteins Hta1, Htb1, Hht2, and Hhf2 from the 500 mM NaCl IP-MS screen (IP vs complement), row clustered.

**Figure S6.**
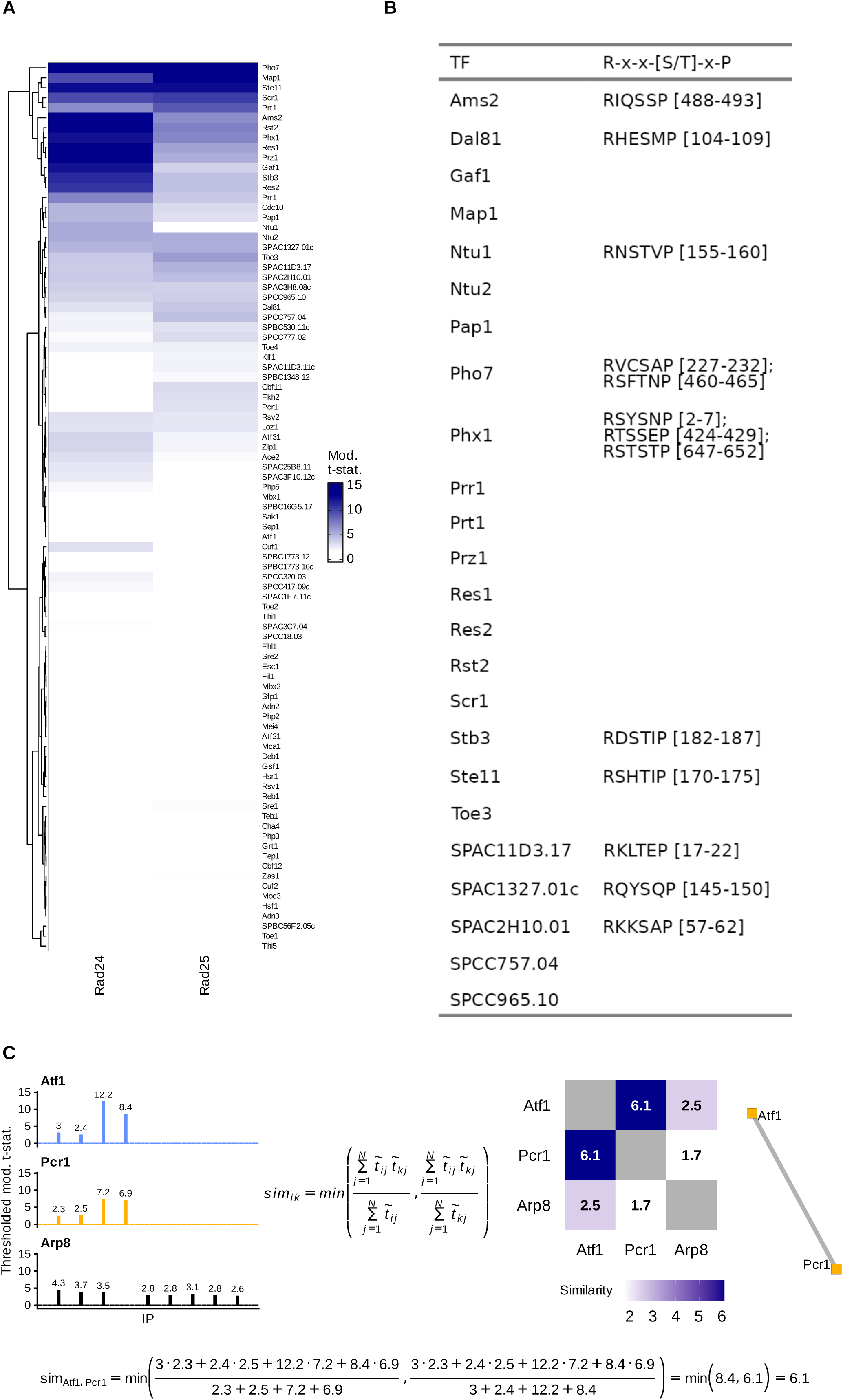
Extensive interactions between TFs and Rad24/Rad25 suggest widespread TF regulation by 14-3-3 proteins, related to Figure 6. (A) Heatmap of moderated t-statistics for Rad24 and Rad25 from the 150 mM NaCl IP-MS screen (IP vs untagged control), row clustered. (B) Table annotating protein sequence motifs matching the pattern (R-x-x-[S/T]-x-P) of TFs interacting with at least one 14-3-3 protein (go to [File](https://github.com/fmicompbio/Spombe_TFome/raw/main/data/FigS6_Rad24_Rad25_interactingTFs_list_motifs.txt)). Amino acid positions of motifs are shown in brackets. (C) Schematic explaining the calculation of pairwise protein similarities across IPs to generate bait-agnostic prey-prey interaction networks using known interacting proteins, Atf1 and Pcr1, alongside an unrelated protein, the INO80 subunit Arp8, as examples. In summary, we truncated the moderated t-statistic profile for each protein across all IPs, setting all negative values or values with an adjusted p-value above 0.1 to 0 and calculated a similarity score for each pair of proteins based on these truncated t-statistics. We computed the inner product of the truncated t-statistic vectors for each two IPs. Next, we divided this number by the sum of the truncated t-statistics to obtain two similarity scores. To avoid overestimating interactions, we selected the minimum of these two scores, resulting in a symmetric matrix. By thresholding the matrix of similarity scores, we create an adjacency matrix, from which we generate a network containing all proteins with at least one remaining edge, with the edge weight indicating the similarity between profiles of two proteins across all IPs. Also see methods.

**Figure S7.**
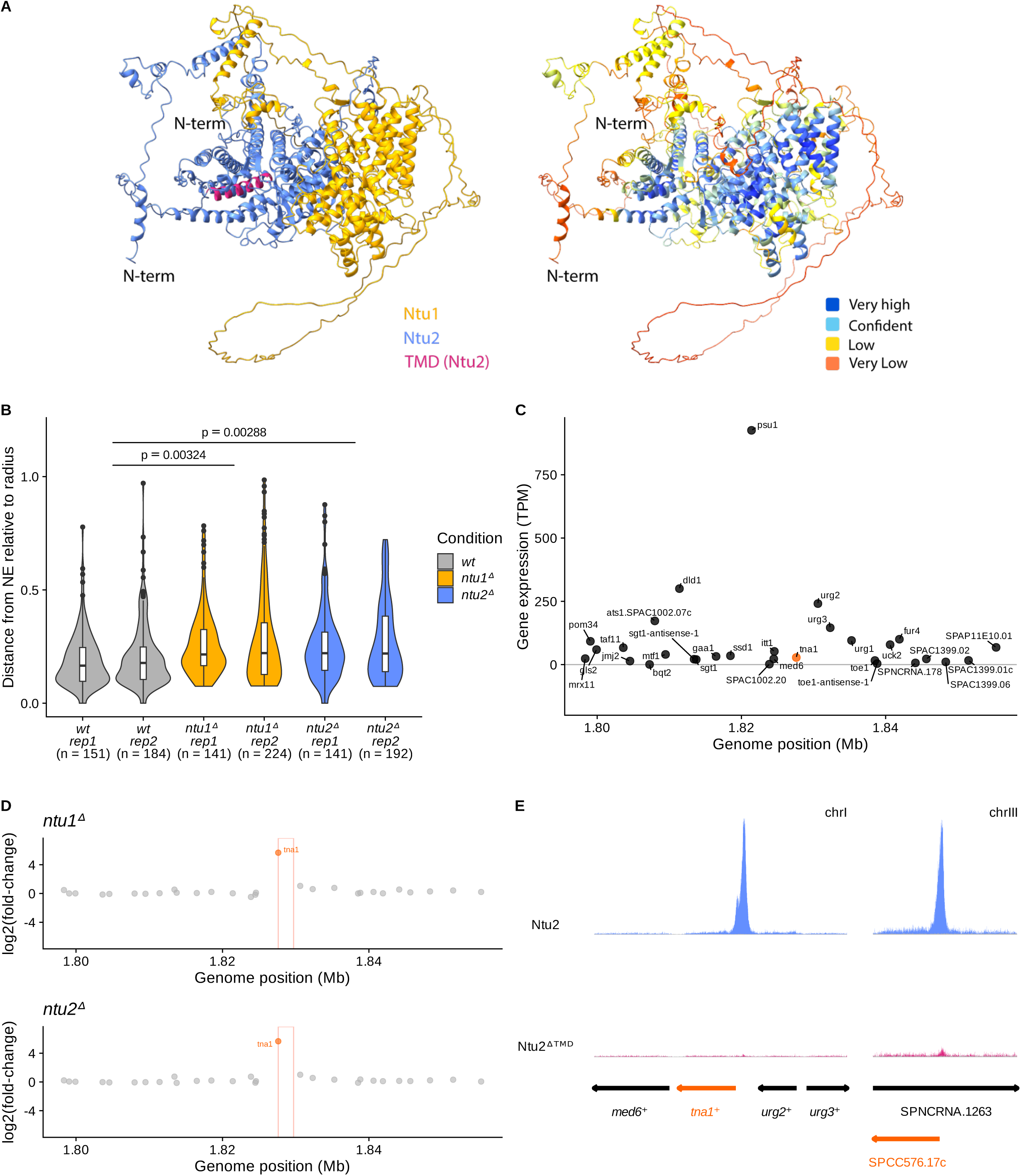
Nattou-mediated gene repression at the nuclear periphery, related to Figure 7. (A) Cartoon models of the Ntu1/Ntu2 interaction (full-length) predicted by AlphaFold^96–98^. Left model: Colored by protein identities with Ntu1 in yellow, Ntu2 in blue, and Ntu2’s predicted TMD in pink. Right model: Colored by model confidence (pLDDT). (B) Quantification of lacI-GFP in *wt*, *ntu1^Δ^*, and *ntu2^Δ^* cells relative to the nuclear envelope (NE). n indicates the number of cells counted in two independent experiments. Statistical analysis was performed by fitting a linear model to the median of the relative distances per replicate, using condition as the sole predictor. (C) mRNA expression levels (wild-type) for genes around the *tna1^+^* locus (30 kb window) in TPMs. (D) Log2(fold-change) gene expression around the *tna1^+^* locus (30 kb window) in *ntu1^Δ^* or *ntu2^Δ^* cells compared to wild type. *tna1^+^* gene indicated by orange box. (E) Genome browser views at the *tna1^+^* (chrI) and SPCC576.17c (chrIII) loci showing relative ChIP-seq fragment densities for Ntu2 (blue) and Ntu2^ΔTMD^ (pink), with input overlay in grey. Scale: CPM per bp, subset to the loci of interest and scaled by dividing by the largest CPM value, separately for each of the two displayed loci.

