## Supplementary material for "A comprehensive *Schizosaccharomyces pombe* atlas of physical transcription factor interactions with proteins and chromatin": Key Resources Table

| REAGENT or RESOURCE | SOURCE | IDENTIFIER |
|---|---|---|
| Antibodies | | |
| Mouse monoclonal anti-FLAG M2 | Sigma-Aldrich | Cat#F1804; RRID: AB_262044 |
| Bacterial and virus strains | | |
| Biological samples | | |
| Chemicals, peptides, and recombinant proteins | | |
| HALT Protease Inhibitor Cocktail (100x) | Thermo Fisher Scientific | Cat#78438 |
| NEBNext® High-Fidelity 2X PCR Master Mix | NEB | Cat#M0541 |
| Dynabeads Protein G | Thermo Fisher Scientific | Cat#10003D |
| Lysyl Endopeptidase (Lys-C) (mass spectrometry grade) | Wako | Cat#125-05061 |
| Trypsin Protease (mass spectrometry grade) | Thermo Fisher Scientific | Cat#90058 |
| Lectin (from *Glycine max*) | Sigma-Aldrich | L1395-5MG |
| Critical commercial assays | | |
| NEBNext Ultra II DNA Library Prep Kit | NEB | E7645 |
| NEBnext Multiplex Oligos for Illumina (Unique dual Index primer pairs) | NEB | E6440, E6442 |
| Illumina Stranded mRNA Prep | Illumina | 20040534 |
| Qubit dsDNA High Sensitivity | Thermo Fisher Scientific | Q32854 |
| Qubit RNA Broad Range | Thermo Fisher Scientific | Q10210 |
| NovaSeq 6000 SP/S1 Reagent kit v1.5 (100 cycles) | Illumina | 20028401/20028319 |
| High Sensitivity D1000 ScreenTape Reagent kit | Agilent | 5067-5585 |
| RNA ScreenTape Reagent kit | Agilent | 5067-5576 |
| HS NGS Fragment Kit (1-6000 bp) | Agilent | DNF-474-1000 |
| MasterPure Yeast RNA Purification Kit | Lucigen | MPY03100 |
| Deposited data | | |
| Chromatin immunoprecipitation sequencing data (ChIP-seq) | GEO | GSE274238 |
| mRNA-sequencing data (mRNA-seq) | GEO | GSE274240 |
| Immunoprecipitation-mass spectrometry data (IP-MS) | PRIDE | PXD054070 |
| ATAC-seq (Existing, public data)[104] | GEO | GSE66386 (GSM1621334-GSM1621338) |

| | | |
|---|---|---|
| ChIP-seq H3 and H3K14ac (Existing, public data)[105] | GEO | GSE108668 (GSM2909801, GSM2909803, GSM2909806, GSM2909807, GSM2909810, GSM2909811) |
| ChIP-seq H3K9me2 and H3K9me3 (Existing, public data)[106] | GEO | GSE182250 (GSM5955195-GSM5955200) |
| GitHub repository archive of code, metadata, and any additional information required to reanalyze the data reported in this paper including fully reproducible reports for all figures | Zenodo | https://zenodo.org/doi/10.5281/zenodo.13270428 |
| Experimental models: Cell lines | | |
| *S. pombe*: Haploid Deletion Library[134] v5.0 | Bioneer | Cat#M-5030H-LT |
| *S. pombe*: *h90 ade6-M216 leu1-32 lys1-131 ura4-D18 his7⁺::lacI-GFP A25::lacOP-ura4⁺-kanr gcut11:mCherry:HygR* | NBRP Japan (https://nbrp.jp/en/) | FY38775[112] |
| *S. pombe*: endogenously anti-FLAG tagged TF library | This paper; available at NBRP Japan (https://nbrp.jp/en/) | FY49691-FY49780 |
| Please see Table S1 for a list of all *S. pombe* strains used in this study | This paper | N/A |
| Experimental models: Organisms/strains | | |
| Oligonucleotides | | |
| Please see Table S2 for a list of primers used in this study | This paper | N/A |
| Recombinant DNA | | |
| Please see Data S1 for all plasmids used in this study | This paper | N/A |
| Software and algorithms | | |
| R v4.3.2 | R Core Team | https://www.r-project.org/ |
| Bioconductor[135] v3.18 | Bioconductor Core Team | https://www.bioconductor.org/ |
| einprot v0.9.3 | Soneson *et al.*[108] | https://github.com/fmicompbio/einprot, commit 27f623ea |
| Bcl2fastq2 v2.20 | Illumina | N/A |
| Real-Time Analysis RTA v3.4.4 | Illumina | N/A |
| FiJi/ImageJ software | Schindelin *et al.*[113] | https://imagej.net/ |
| ScanProsite tool | De Castro *et al.*[93] | https://prosite.expasy.org/scanprosite/ |
| NCBI Conserved Domain (CD) Search | Wang *et al.*[17] | https://www.ncbi.nlm.nih.gov/Structure/cdd/wrpsb.cgi |

| | | |
|---|---|---|
| AlphaFold2 and AlphaFold-Multimer | Jumper *et al.*[96]<br>Varadi *et al.*[97]<br>Evans *et al.*[98] | https://github.com/google-deepmind/alphafold |
| GUIFold | Kempf and Cavadini[116] | https://github.com/fmi-basel/GUIFold |
| ChimeraX v1.6.1 | Pettersen *et al.*[117] | https://www.rbvi.ucsf.edu/chimerax |
| LinkedCharts | Ovchinnikova and Anders[114] | https://anders-biostat.github.io/linked-charts/ |
| IGV for interactive genome visualization | Robinson *et al.*[115] | https://github.com/igvteam/igv.js |
| tximeta v1.16.1 | Love *et al.*[130] | https://doi.org/doi:10.18129/B9.bioc.tximeta |
| fishpond v2.4.1 | Zhu *et al.*[131] | https://doi.org/doi:10.18129/B9.bioc.fishpond |
| DESeq2 v1.42.1 | Love *et al.*[132] | https://doi.org/doi:10.18129/B9.bioc.DESeq2 |
| STAR v2.7.10b | Dobin *et al.*[128] | https://github.com/alexdobin/STAR |
| Salmon v1.9.0 | Patro *et al.*[126] | https://github.com/COMBINE-lab/salmon |
| DeepTools v3.3.1 | Ramírez *et al.*[129] | https://github.com/deeptools/deepTools/ |
| MaxQuant v2.2.0.0 | Cox and Mann[118] | https://www.maxquant.org/ |
| scater v1.26.1 | McCarthy *et al.*[120] | https://doi.org/doi:10.18129/B9.bioc.scater |
| limma v3.54.2 | Ritchie *et al.*[20] | https://doi.org/doi:10.18129/B9.bioc.limma |
| rtracklayer v1.58.0 | Lawrence *et al.*[133] | https://doi.org/doi:10.18129/B9.bioc.rtracklayer |
| ggplot2 v3.5.0 | Wickham[136] | https://doi.org/10.32614/CRAN.package.ggplot2 |
| ComplexHeatmap v2.18.0 | Gu *et al.*[137] | https://doi.org/doi:10.18129/B9.bioc.ComplexHeatmap |
| QuasR v1.38.0 | Gaidatzis *et al.*[121] | https://doi.org/doi:10.18129/B9.bioc.QuasR |
| cutadapt v3.7 | Martin[122] | https://github.com/marcelm/cutadapt/ |

| | | |
|---|---|---|
| MACS2 v2.2.7.1 | Zhang *et al.*[123] | https://github.com/macs3-project/MACS |
| MEME v5.5.2 | Bailey[124] | https://github.com/cinquin/MEME |
| universalmotif v1.16.0 | Tremblay and Nystrom[125] | https://doi.org/10.18129/B9.bioc.universalmotif |
| imputeLCMD v2.1 | Lazar *et al.*[119] | https://github.com/cran/imputeLCMD |
| Other | | |
| TFexplorer webtool | This paper | https://data.fmi.ch/TFexplorer/ |
| Code, metadata, and any additional information required to reanalyze the data reported in this paper including fully reproducible reports for all figures | This paper | GitHub: https://github.com/fmicompbio/Spombe_TFome Zenodo: https://zenodo.org/doi/10.5281/zenodo.13270428 |
| FastPrep-24 5G bead beating grinder | MP Biomedicals | Cat#6005500 |
| MagnaBot® II Magnetic Separation Device | Promega | Cat#V8351 |
| DynaMag™-2 Magnet | Thermo Fisher Scientific | Cat#12321D |
| Bioruptor® Pico | Diagenode | Cat#B01060010 |
| MatTek glass bottom dish | MatTek | P35G-1.5-10-C |
